# Semenogelin-1 Inhibition of Mouse Sperm Hyperactivation Reveals Two Functional Domains Modulating CatSper Channel

**DOI:** 10.1101/2025.09.05.674523

**Authors:** Noemia A. P. Mariani, Natália C. M. Santos, Juliana J. Andrade, Alexandre D. Andrade, Beatriz Rezende, Tomás J. Steeman, Analia G. Novero, José C. de Lima-Junior, Giulia Calderaro, Ana Clara F. Themer, Hélio Kushima, Mariano G. Buffone, Darío Krapf, Polina V. Lishko, Erick J. R. Silva

**Affiliations:** Department of Biophysics and Pharmacology, Biosciences Institute of Botucatu, São Paulo State University, Botucatu, SP, Brazil; Institute of Molecular and Cellular Biology of Rosario, Rosario, SF, Argentina; Department of Cell Biology and Physiology, Washington University in St. Louis, School of Medicine, St. Louis, MO, United States; Center for the Investigation of Membrane Excitability Diseases (CIMED), WashU Medicine, St. Louis, MO, United States; Institute of Biology and Experimental Medicine, Buenos Aires, BA, Argentina

**Keywords:** Sperm motility, Epididymal protease inhibitor, Spermatozoon, Ion Channels, Seminal Plasma, Semenogelins

## Abstract

Seminal plasma is essential for sperm survival and function after ejaculation. Semenogelin-1 (SEMG1), the predominant seminal plasma protein, transiently suppresses sperm motility and hyperactivation after ejaculation through epididymal protease inhibitor (EPPIN) binding. However, the molecular mechanism underlying SEMG1-mediated inhibition of hyperactivation remains unclear. Here, we tested the hypothesis that SEMG1 inhibits CatSper, a sperm-specific Ca^2+^ channel crucial for hyperactivation. Full-length recombinant mouse SEMG1 (mSEMG1; Q32-G375) inhibited both progressive motility and hyperactivation; the latter was not recovered with NH_4_Cl-induced alkalinization, indicating an effect downstream of capacitation-associated intracellular alkalinization. Electrophysiological recordings revealed that mSEMG1 reduced CatSper currents at physiologically relevant concentrations. Truncated mSEMG1 fragments mSEMG1^Q32-V118^ and mSEMG1^R98-G375^, but not mSEMG1^Y221-G375^, inhibited sperm hyperactivation and CatSper currents to a similar extent as full-length mSEMG1. Notably, only mSEMG1^R98-G375^ retained full EPPIN-binding capacity. Together, our findings identify two functional domains within mSEMG1 (Q32-V118 and R98-S220) that inhibit sperm hyperactivation, suppress CatSper currents and differ in their EPPIN-binding capacities. These functional domains represent promising prototypes for the design of spermiostatic molecules, offering additional avenues for non-hormonal male contraception.

**Summary statement:** We report that mouse SEMG1 contains two distinct functional domains that inhibit sperm hyperactivation and reduce CatSper currents, with implications for non-hormonal male contraception.

## Introduction

To successfully fertilize the egg, spermatozoa must undergo their final maturation within the female reproductive tract, a process known as capacitation (Chang, 1951; Austin, 1952; Stival et al., 2016; Freitas et al., 2017). Sperm capacitation entails a complex cascade of molecular events, including cholesterol efflux from the plasma membrane, tyrosine phosphorylation of sperm proteins, increases in intracellular pH (pH_i_) and Ca^2+^ concentrations ([Ca^2+^]_i_), as well as membrane hyperpolarization. Sperm capacitation allows spermatozoa to undergo acrosome exocytosis in response to appropriate stimuli and to transition from progressive motility to a vigorous, asymmetric, whip-like flagellar motion known as hyperactivation (López-González et al., 2014; Freitas et al., 2017; Ritagliati et al., 2018; Vyklicka and Lishko, 2020).

Among the molecular pathways involved in sperm hyperactivation, the rise in intracellular Ca^2+^ levels is critical (Darszon et al., 2011). Although multiple mechanisms promote Ca^2+^ influx in spermatozoa, the activation of the voltage-dependent and alkalinization-activated Ca^2+^ channel CatSper is considered the most relevant (Lishko and Mannowetz, 2018). The CatSper channel consists of a complex of at least fourteen proteins: four pore-forming subunits (CatSper1–4) and 10 accessory subunits (CatSper β, γ, δ, ε, ζ, τ, η, θ, EFCAB9, and transporter SLCO6C1) (Lishko and Mannowetz, 2018; Hwang et al., 2019; Lin et al., 2021). This large multiprotein complex, known as the CatSpermasome, is arranged as four longitudinal lines spanning the entire length of the sperm principal piece, facilitating structural and functional coupling between CatSper complexes and triggering powerful Ca^2+^ waves along the sperm flagellum (Bystroff, 2018; Vyklicka and Lishko, 2020; Lin et al., 2021; Zhao et al., 2022). Notably, CatSper deficiency in men and rodents leads to infertility due to a failure of sperm hyperactivation, while no other phenotypical abnormalities are observed (Carlson et al., 2005; Avenarius et al., 2009; Hildebrand et al., 2010; Chung et al., 2017).

While sperm hyperactivation is crucial for fertilization, it must remain tightly regulated until spermatozoa reach the oviduct (De Jonge, 2017). In this context, seminal plasma components play critical roles in modulating sperm hyperactivation and other capacitation-associated events (Vickram et al., 2022). Semenogelin-1 (SEMG1), the predominant protein in seminal plasma, is secreted by the seminal vesicles and incorporated onto the sperm surface during ejaculation through its interaction with epididymal protease inhibitor (EPPIN) in both humans and rodents (Wang et al., 2005; Silva et al., 2012; Silva et al., 2013; Mariani et al., 2020). SEMG1 binds to freshly ejaculated spermatozoa, acting as a decapacitation factor and blocking sperm motility, hyperactivation, and acrosome exocytosis (Kawano and Yoshida, 2007; de Lamirande and Lamothe, 2010; Mitra et al., 2010; Araki et al., 2015). In humans, this inhibitory effect is maintained until the serine protease prostate-specific antigen (PSA) cleaves SEMG1, resulting in semen liquefaction and activation of sperm motility (Robert and Gagnon, 1999). Furthermore, SEMG1 promotes sperm survival in the uterus, as evidenced by the low viability of spermatozoa from *Semg1*^-/-^ mice collected from the female reproductive tract during natural mating, causing a severe subfertility phenotype (Kawano et al., 2014). Together, these findings indicate that SEMG1 contributes to the regulation of sperm function and metabolism during their journey through the female reproductive tract. However, the precise molecular mechanisms underlying SEMG1’s roles and the contribution of EPPIN binding to these effects remain poorly understood.

Initial studies have shown inhibitory effects of SEMG1 on sperm pH_i_ and [Ca^2+^]_i_ (O’Rand and Widgren, 2012). Building upon these findings, we hypothesize that SEMG1 inhibits CatSper activity, thereby altering Ca^2+^ homeostasis and downstream pathways required for sperm hyperactivation. To test this hypothesis, we investigated the effects of recombinant mouse SEMG1 (mSEMG1) on sperm motility, hyperactivation, and CatSper currents. We further investigated the contribution of distinct SEMG1 domains and assessed the relevance of EPPIN binding in mediating their effects. Our results revealed that mSEMG1 inhibits hyperactivation by modulating CatSper currents, an effect mediated by distinct domains within the Q32-V118 and R98-S220 sequences. Remarkably, only the R98-S220 sequence bound EPPIN to the full extent, indicating that SEMG1 regulates sperm hyperactivation through domain-specific mechanisms associated with differential EPPIN interaction. Our findings provide novel mechanistic insights into how mSEMG1 prevents premature sperm hyperactivation and open avenues for exploring its functional domains as templates for the rational design of ligands with male contraceptive applications targeting spermatozoa.

## Results

### Mouse SEMG1 inhibits sperm progressive and hyperactivated motilities

The sperm motility pattern changes from progressive to hyperactivated upon capacitation (Katz and Yanagimachi, 1980; Suárez and Osman, 1987). Previous studies have shown that both human and mouse SEMG1 can regulate sperm function, including motility, after ejaculation (Mitra et al., 2010; Silva et al., 2013; Sakaguchi et al., 2020). Consistent with these observations, we observed that SEMG1 was not detected in mouse spermatozoa isolated from the cauda epididymidis, but became detected following incubation with seminal vesicle fluid (Fig. S1). To gain insights into the effects of SEMG1 on sperm swimming behavior, we incubated murine spermatozoa isolated from the cauda epididymidis in capacitating medium, in either the absence (control) or in the presence of increasing concentrations of full-length recombinant mouse SEMG1 (mSEMG1; Q32-G375). mSEMG1 reduced sperm motility only at 25 µM (∼25% inhibition) after 20 min (p = 0.0005) and 120 min (p = 0.011) of incubation compared to the control (Fig. 1A, Fig. S2, Video S1). Concurrently, mSEMG1 increased the percentage of slow-motile spermatozoa at 10 µM (p = 0.026) and 25 µM (p = 0.001) after 20 min of incubation compared to the control (Fig. 1B, Fig. S2). mSEMG1 did not affect sperm viability at the concentrations tested, confirming that its effects on sperm motility were not associated with sperm death (Fig. 1C).

**Figure 1.**
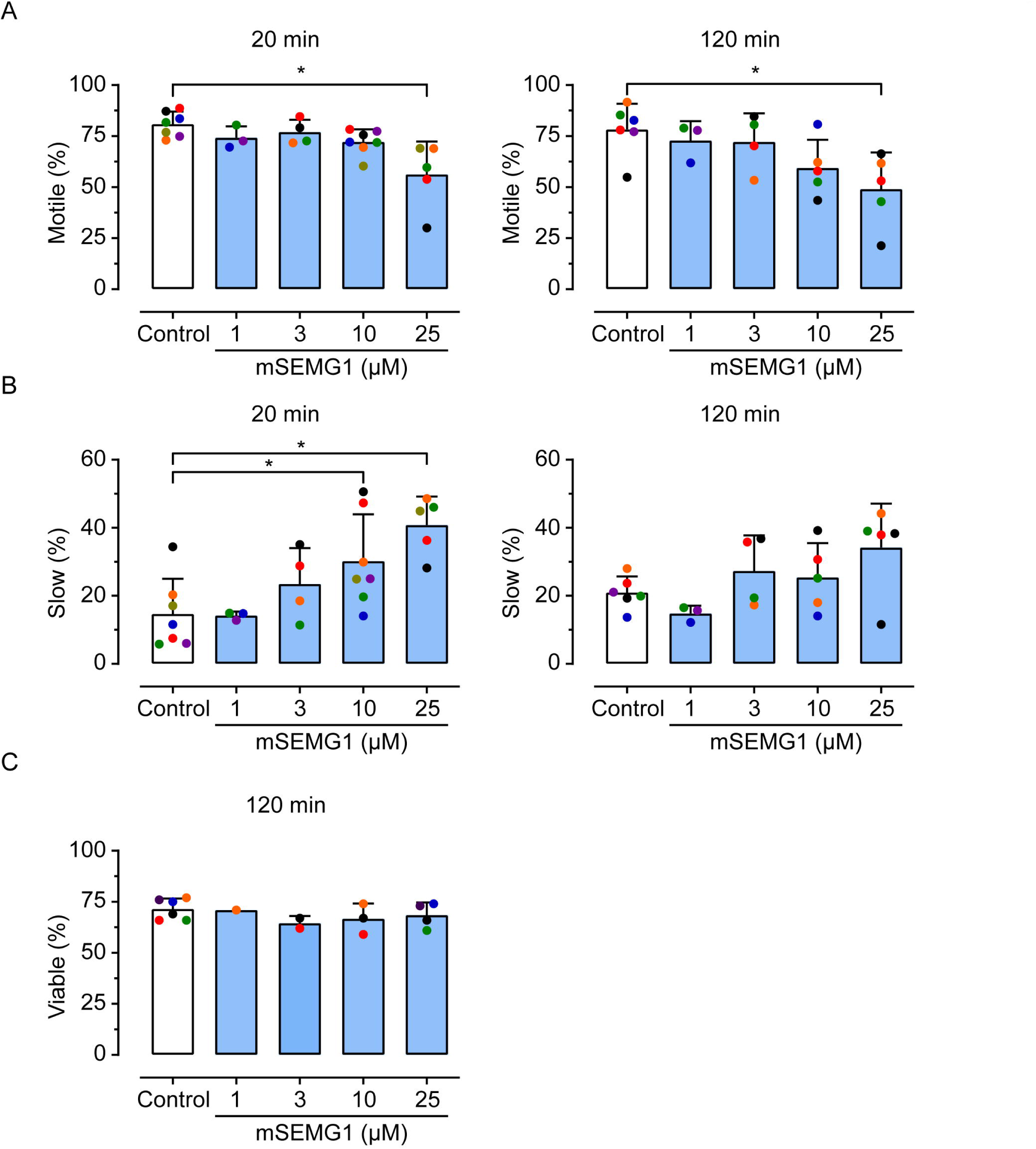
Effects of mSEMG1 on murine sperm motility and viability. (A and B) Percentage of total and slow motility of spermatozoa after incubation with mSEMG1. (C) Percentage of viable spermatozoa after incubation with mSEMG1. Spermatozoa were incubated in capacitating HTF medium in the absence (control) or presence of increasing concentrations of mSEMG1 (1, 3, 10, and 25 μM). Sperm motility was assessed after 20 and 120 min of capacitation, while sperm viability was assessed after 120 min. Dots with the same color represent the values from an independent experiment. Asterisks indicate statistically significant differences from the control group (p<0.05, ANOVA followed by Dunnett’s test). Percentage data were transformed using arcsine square-root values before statistical analysis. Results are presented as mean ± SD values from independent experiments using sperm samples from 3-7 mice.

mSEMG1 displayed a more potent inhibitory effect on hyperactivated motility by inducing a concentration-dependent inhibition with an IC_50_ of 5.2 µM (95% confidence interval; 95% CI 1.3–21.2 µM). The maximum inhibition of ∼90% was reached at 25 µM (p = 0.0003) after 120 min, compared to the control (Fig. 2A, B; Fig. S2). Consistently, 25 µM mSEMG1 decreased curvilinear velocity (VCL, p = 0.006) and amplitude of lateral head displacement (ALH, p = 0.005), kinematic parameters associated with vigorous sperm movements (Cancel et al., 2000) (Fig. 2C). Notably, mSEMG1 effects on hyperactivation were not rescued by intracellular alkalization induced by NH_4_Cl (Fig. 2D), a condition that enhance sperm hyperactivation via alkalinization evoked activation of CatSper (Ren et al., 2001; Kirichok et al., 2006) (Fig. 2D). Similarly, mSEMG1 inhibited progressive motility (IC_50_ 10.1 µM, 95% CI 2.6–38.5 µM), and decreased both average path velocity (VAP) and straight-line velocity (VSL), kinematic parameters associated with sperm progressiveness (Fig. S2). Altogether, these results indicate that mSEMG1 inhibits mouse sperm progressive motility and hyperactivation by modulating signaling pathways downstream of intracellular alkalinization.

**Figure 2.**
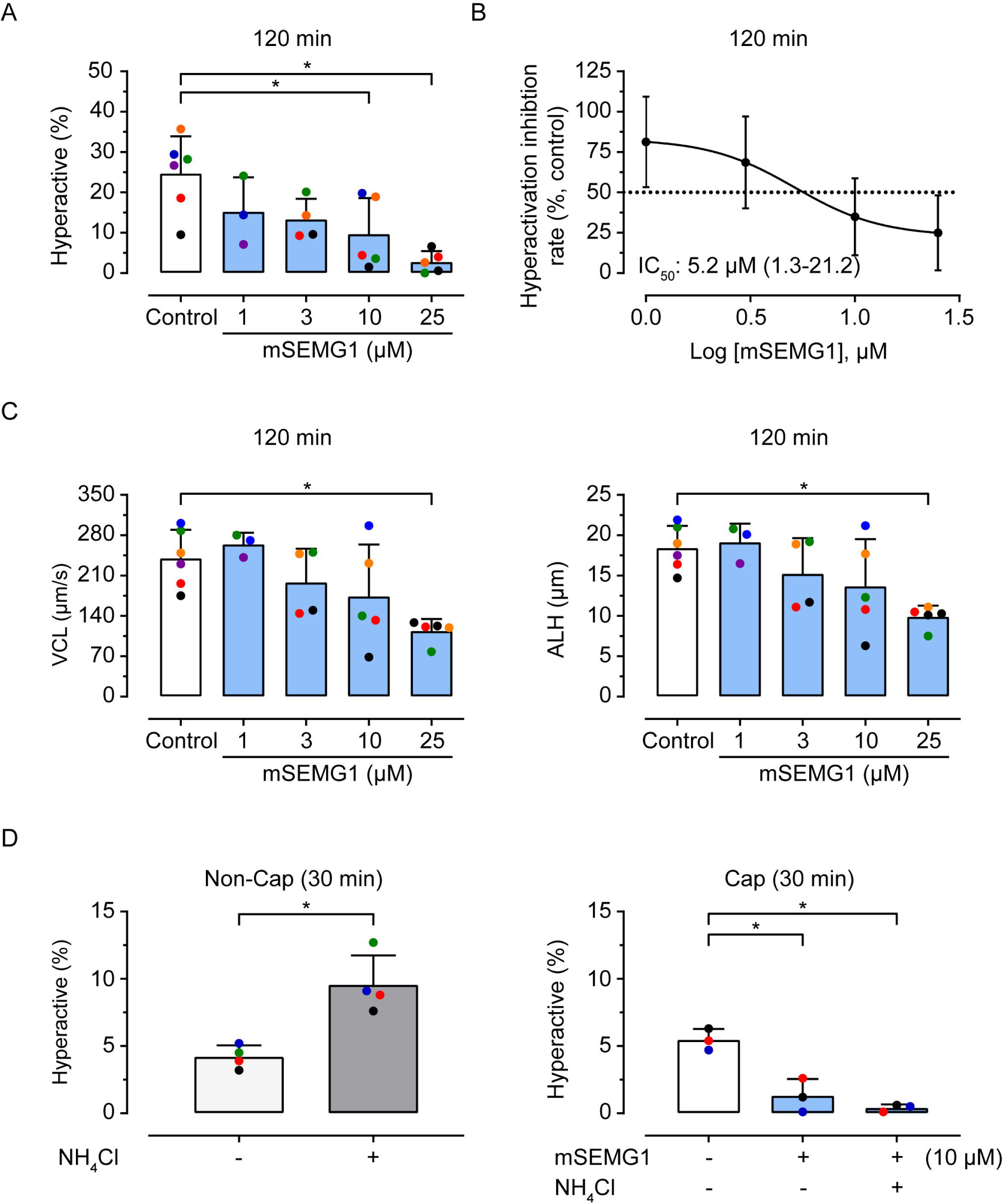
Effects of mSEMG1 on hyperactivated motility. (A) Percentage of hyperactivated motility of spermatozoa after incubation with mSEMG1. (B) Concentration-response curve of hyperactivation inhibition rate after incubation with mSEMG1. Data were expressed as mean percentage values relative to the control group, calculated as (% hyperactivated spermatozoa in the treated sample divided by % hyperactivated spermatozoa in the matched control) × 100. The half-maximal Inhibitory concentration (IC_50_) and its 95% confidence interval) are presented. (C) Curvilinear velocity (VCL) (left) and amplitude of lateral head displacement (ALH) (right) of spermatozoa after incubation with mSEMG1. Spermatozoa were isolated from the cauda epididymidis and incubated in capacitating HTF medium in the absence (control) or presence of increasing concentrations of mSEMG1 (1, 3, 10, and 25 μM). (D) Percentage of hyperactivated spermatozoa in the presence or absence of NH_4_Cl. Cells were incubated in non-capacitating medium (Non-Cap), supplemented or not with NH_4_Cl (left), or in capacitating medium (Cap), supplemented or not with NH_4_Cl in the absence or presence of 10 µM mSEMG1 (right). Sperm motility was assessed after 30 and 120 min of capacitation. Dots with the same color represent the values from an independent experiment. Asterisks indicate statistically significant differences from the control group (p<0.05, ANOVA followed by Dunnett’s test). Percentage data were transformed using arcsine square-root values before statistical analysis. Results are presented as mean ± SD values from independent experiments using sperm samples from 3-6 mice.

### Mouse SEMG1 is a CatSper channel inhibitor

Activation of the CatSper channel leads to a rise in intracellular Ca^2+^ concentration, which is essential for the development of hyperactivated motility in both murine and human spermatozoa (Quill et al., 2003; Carlson et al., 2005; Young et al., 2024). Thus, we explored whether mSEMG1 inhibits hyperactivation by affecting the CatSper channel. To do so, we conducted whole-cell patch clamp experiments using non-capacitated murine spermatozoa to record CatSper currents (I*_CatSper_*) before and after mSEMG1 treatment.

I*_CatSper_* were elicited by voltage ramps from -80 to +80 mV from a holding potential of 0 mV (Fig. 3A). Under these conditions, no CatSper current was detected in a high-saline (HS) extracellular bath solution containing Ca^2+^ and Mg^2+^ (Table S1), which represents the baseline recording (Fig. 3B). The remaining current under this condition was attributed to a leak current and was negligible (mean ± SD: −16.58 ± 9.59 pA; n = 21). Upon perfusion with divalent-free Cs^+^-based (DVF) bath solution (Table S2), the characteristic monovalent CatSper current was recorded, as previously described (Kirichok and Lishko, 2011; Liu et al., 2021) (Fig. 3B). Following the DVF perfusion, sperm cells were exposed to 5 µM mSEMG1, which suppressed both inward and outward CatSper currents (Fig. 3B, C). After subtracting the leak currents, the averaged (± standard deviation, SD) control inward I*_CatSper_* in DVF at - 80 mV was 218.4 ± 62.3 pA (Fig. 3B). Under the same conditions, exposure to 5 µM mSEMG1 inhibited the inward I*_CatSper_*to 150.8 ± 46.9 pA (∼31% suppression; p = 0.019) (Fig. 3B). No currents were detected when spermatozoa from *CatSper1*^-/-^ mice were tested either in the absence or presence of mSEMG1 (Fig. 3B, insert). mSEMG1 inhibition was reversible, as CatSper conductance recovered upon continuous washing in DVF (177.0 ± 24.3 pA; p = 0.378) (Fig. 3B).

**Figure 3.**
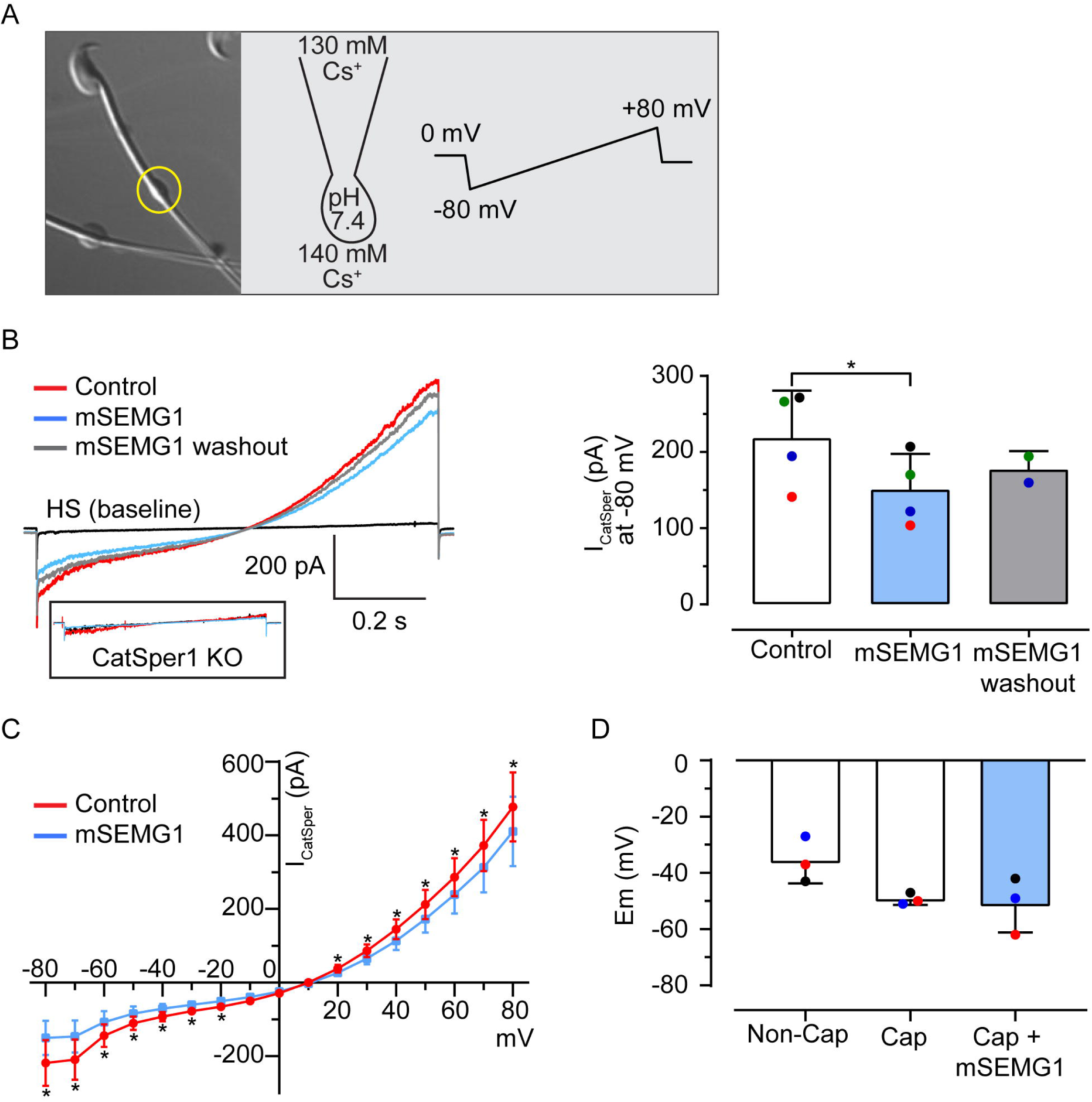
Effects of mSEMG1 on I*_CatSper_*. (A) Representative DIC image of a murine spermatozoon with cytoplasmic droplet (yellow circle) used for patch-clamp recording. Right: Cs⁺ concentration and pH of the pipette and bath solutions are indicated. Voltage ramps (-80 mV to 80 mV, 1 s) were applied in HS and in DVF to record I*_CatSper_*. Baseline currents were recorded in HS (black), which lacked I*_CatSper_* due to its inhibition by extracellular magnesium. (B) Left: I*_CatSper_* recordings in response to voltage ramps in DVF before (control) (red) and after addition of 5 μM mSEMG1 (blue), including washout (grey), and recorded from wild-type and CatSper1-KO (inset) spermatozoa. Traces represent recordings from the same spermatozoon. Right: I*_CatSper_*recordings (mean ± SD; pA) obtained at −80 mV in DVF solution before and after exposure to 5 μM mSEMG1. I*_CatSper_*recording was repeated in DVF after mSEMG1 was washed out. Sperm cell capacitance ranged from 2.0 to 2.6 pF. Dots with the same color represent current values from the same cell. Results are presented as mean ± SD from 4 spermatozoa isolated from 4 mice, and from 2 of these spermatozoa for the washout. (C) Current-voltage (I-V) relationships were calculated from amplitudes shown in panel A in the DVF solution before and after 5 μM mSEMG1 treatment. (D) Quantification of membrane potential (Em) in non-capacitated, capacitated, and capacitated spermatozoa treated with 10 μM mSEMG1. Membrane potential was assessed after 30 min of capacitation. Dots with the same color represent values from independent experiments. Asterisks indicate statistically significant differences from the control group (p < 0.05, ANOVA followed by Dunnett’s test or paired t-test). Results are presented as mean ± SD from independent experiments using sperm samples from 3 mice.

In addition to intracellular alkalization, CatSper is sensitive to membrane voltage (Kirichok et al., 2006; Vyklicka and Lishko, 2020). During capacitation of mouse spermatozoa, intracellular pH increases, activating the sperm-specific, pH-sensitive K⁺ channel SLO3 and promoting membrane hyperpolarization through K⁺ efflux (Schreiber et al., 1998; Zeng et al., 2011; Zeng et al., 2015). Because murine SLO3 is highly sensitive to intracellular alkalinization, sperm membrane potential (Em) also provides an indirect readout of pHi. We therefore asked whether mSEMG1 altered Em during capacitation. As expected, capacitated spermatozoa displayed a more negative membrane potential in comparison to non-capacitated spermatozoa (Fig. 3D, Fig. S3). Capacitation-induced membrane hyperpolarization was not affected by 10 µM mSEMG1 (Fig. 3D, Fig. S3). These results indicate that mSEMG1 inhibits CatSper activity at low micromolar concentrations comparable to those required to inhibit sperm hyperactivation, without affecting the Em. Since Em reflects the net activity of all conductances and electrogenic transport processes, and is dominated by K⁺ conductance in capacitated mouse spermatozoa, this observation neither excludes an effect of mSEMG1 on other conductances nor establishes a direct action on CatSper (see Discussion)

### Mouse SEMG1 Q32-V118 and R98-S220 sequences independently contribute to the inhibition of sperm hyperactivation and CatSper currents

Despite the low conservation in their primary sequences, both human and mouse SEMG1 exhibit a single cysteine residue (C239 in human SEMG1 and C97 in mouse SEMG1), as well as internal repetitive amino acid sequences. Downstream of the cysteine residue, mouse SEMG1 contains two 31-amino acid repeats (K119-S149 and K177-S207) and multiple short repeats of two (KS), three (GGS, which is also present in human SEMG1), or six-eight (QVKSSVSQ or QVKSSGSQ) amino acid residues that become less frequent toward its C-terminal region (Fig. 4A). Previous studies demonstrated that sequences within human SEMG1 that flank the cysteine residue are essential for its interaction with spermatozoa, EPPIN binding, and modulation of sperm motility (Mitra et al., 2010; Silva et al., 2013); however, the sequences containing the long repeats (K282-E319 and K342-E379) were not specifically evaluated. Based on these observations, we obtained three truncated mSEMG1 constructs (i) mSEMG1^Q32-V118^, which contains its unique cysteine residue, (ii) mSEMG1^R98-G375^, which lacks the cysteine residue but includes the two long repeats, and (iii) mSEMG1^Y221-G375^, which lacks both the cysteine residue and the long repeats (Fig. 4A), allowing us to evaluate the relevance of these sequences to the inhibition of sperm hyperactivation and CatSper.

**Figure 4.**
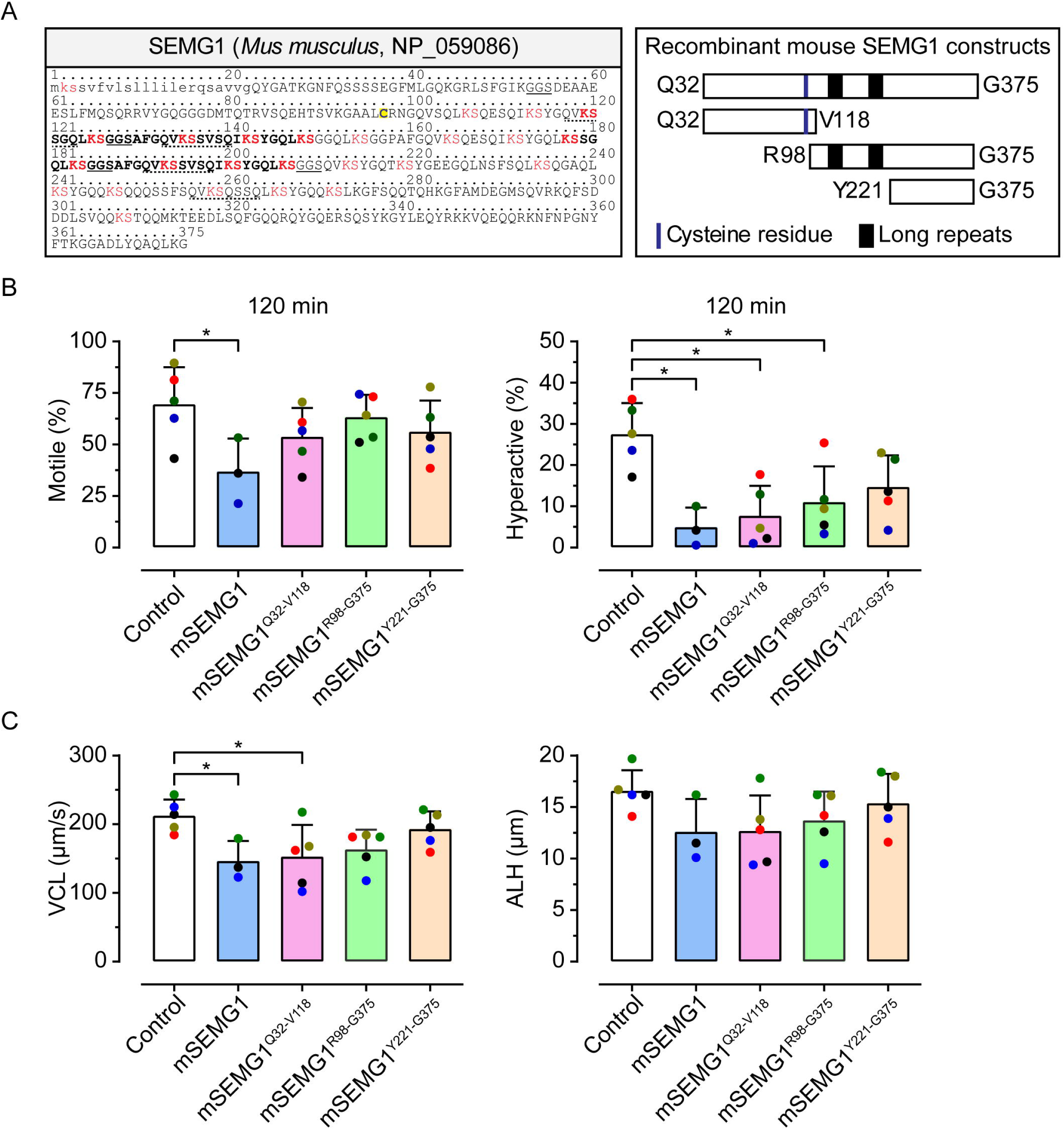
Mouse SEMG1 sequence, recombinant constructs, and their effects on sperm motility. (A) Left: Amino acid sequence of mouse SEMG1. The signal peptide is shown in lowercase letters. Amino acid repeats are indicated in bold. The GGS region is underlined, while the KS region and the single cysteine residue are highlighted in red and yellow, respectively. Right: Schematic representation of mSEMG1 full-length and truncated constructs used in the experiments. The mSEMG1 constructs include a 6x-His tag. (B) Percentage of total motile (left) and hyperactivated (right) spermatozoa after incubation with mSEMG1. (C) Curvilinear velocity (VCL) (left) and amplitude of lateral head displacement (ALH) (right) of spermatozoa after incubation with mSEMG1 constructs. Spermatozoa were isolated from the cauda epididymidis and incubated in capacitating HTF medium in the absence (control) or presence of mSEMG1 constructs (25 μM). Sperm motility was assessed after 120 min. Dots with the same color represent the values from an independent experiment. Asterisks indicate statistically significant differences from the control group (p<0.05, ANOVA followed by Dunnett’s test). Percentage data were transformed using arcsine square-root values before statistical analysis. Results are presented as mean ± SD values from independent experiments using sperm samples from 3-5 mice.

Neither of the truncated mSEMG1 constructs affected sperm motility in a significant manner (Fig. 4B). Nevertheless, mSEMG1^Q32-V118^ and mSEMG1^R98-G375^, but not mSEMG1^Y221-G375^, inhibited hyperactivated motility compared to the control (p = 0.004 and 0.028, respectively) to a similar extent as the full-length mSEMG1 (Fig. 4B, C, Fig. S4, Video S2). mSEMG1^Q32-V118^ also reduced the associated kinematic parameter VCL (p = 0.029), and a similar trend was observed for mSEMG1^R98-G375^ (p = 0.084) (Fig. 4C). Consistent with these findings, both mSEMG1^Q32-V118^ and mSEMG1^R98-G375^ suppressed inward and outward I*_CatSper_* (Fig. 5A, B, Fig. S4). mSEMG1^Y221-G375^ also inhibited CatSper, albeit to a lesser extent than the rest of the truncated constructs (Fig. 5C, Fig. S4). At 5 µM concentration, mSEMG1^Q32-V118^, mSEMG1^R98-G375^, and mSEMG1^Y221-G375^ suppressed the inward I*_CatSper_*at -80 mV to values equivalent to ∼56%, ∼66%, and ∼77% of those observed with their respective controls, corresponding to mean differences of of 93.6 (p = 0.007), 108.6 (p < 0.0001), and 55.9 pA (p = 0.011), respectively (Fig. 5A-C, Fig. S4). Neither of the truncated mSEMG1 constructs affected sperm viability, confirming that its effects on sperm motility were not associated with sperm death (Fig. S4). Although the C-terminal region also displayed a partial inhibitory effect on CatSper currents, its effect on sperm hyperactivation was not significant under our experimental conditions. These results indicate that sequences within the mSEMG1 Q32-V118 and R98-S220 regions, which encompass cysteine and the long repeats, contain domains sufficient to modulate sperm hyperactivation and CatSper.

**Figure 5.**
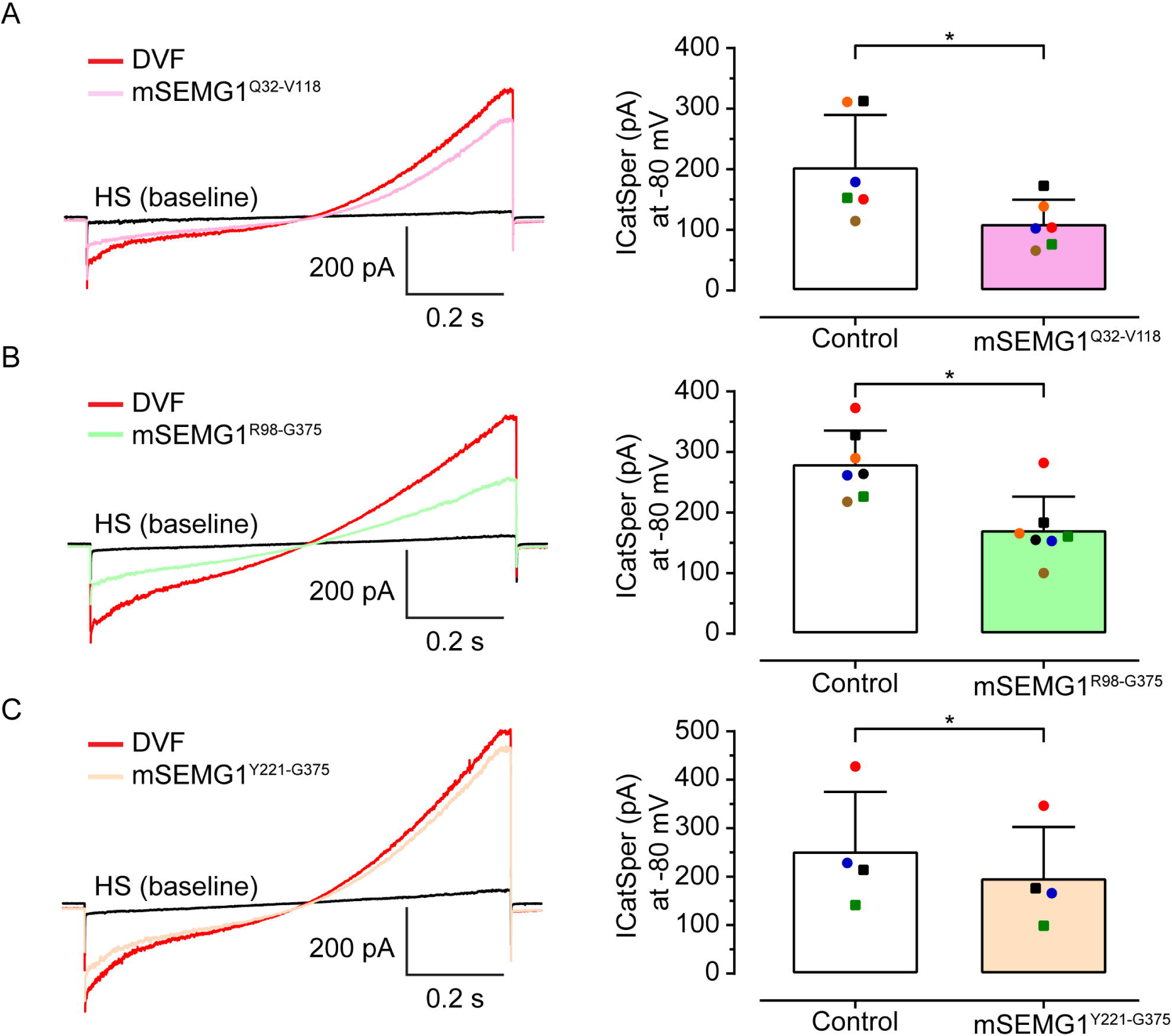
Effects of mSEMG1 fragments on I*_CatSper_*. (A-C) Left: Representative I*_CatSper_*recordings in response to voltage ramps in DVF (control) before (red) and after 5 μM mSEMG1^Q32–V118^ (pink), mSEMG1^R98–G375^ (green), and mSEMG1^Y221–G375^ (orange), recorded from wild-type mouse spermatozoa. Baseline currents are recorded in HS (black). Traces represent recordings from the same spermatozoon. Right: Quantification of I*_CatSper_* (mean ± SD; pA) at −80 mV in DVF solution before and after addition of the respective mSEMG1 fragments. Sperm cell capacitance ranged from 2.1 to 2.8 pF. Dots with the same color represent values from the same cell. Asterisks indicate statistically significant differences from the control group (p < 0.05, paired t-test). Square dots in each panel represent recordings from sperm cells from the same mouse. Results are presented as mean ± SD from 6, 7, and 4 spermatozoa isolated from 5, 6, and 3 mice for panels A, B, and C, respectively.

### Mouse SEMG1 R98-G220 sequence contains the main EPPIN binding domain

Our previous study demonstrated the co-immunoprecipitation and co-localization of SEMG1 and EPPIN in mouse spermatozoa pre-incubated with seminal vesicle fluid, suggesting that EPPIN serves as a docking site for SEMG1 on the mouse sperm surface (Mariani et al., 2020). Here, we confirmed the interaction between mouse EPPIN and SEMG1 using the AlphaScreen (Amplified Luminescent Proximity Homogeneous Assay Screen) assay (Fig. 6A), which has been previously employed to investigate the binding of human EPPIN and SEMG1 (Silva et al., 2012; Silva et al., 2013). Using mSEMG1 and full-length recombinant mouse EPPIN (mEPPIN; L24-T134), we observed a maximum interaction signal peaking at 2 h and remaining stable for at least 8 h (Fig. 6B). The interaction signals showed a concentration-dependent pattern when mSEMG1 and mEPPIN were titrated, with calculated EC_50_ (95% CI) of 23.6 nM (20.0–27.8 nM) and 1.0 nM (0.3–3.3 nM), respectively (Fig. 6C). Interestingly, when recombinant human SEMG1 (hSEMG1, G26-R281) replaced mSEMG1, we observed a concentration-dependent saturable binding curve, with a calculated EC_50_ of 8.3 nM (6.8– 18.5 nM), confirming the conservation of the EPPIN binding sites in human and mouse SEMG1 sequences (Fig. S5).

**Figure 6.**
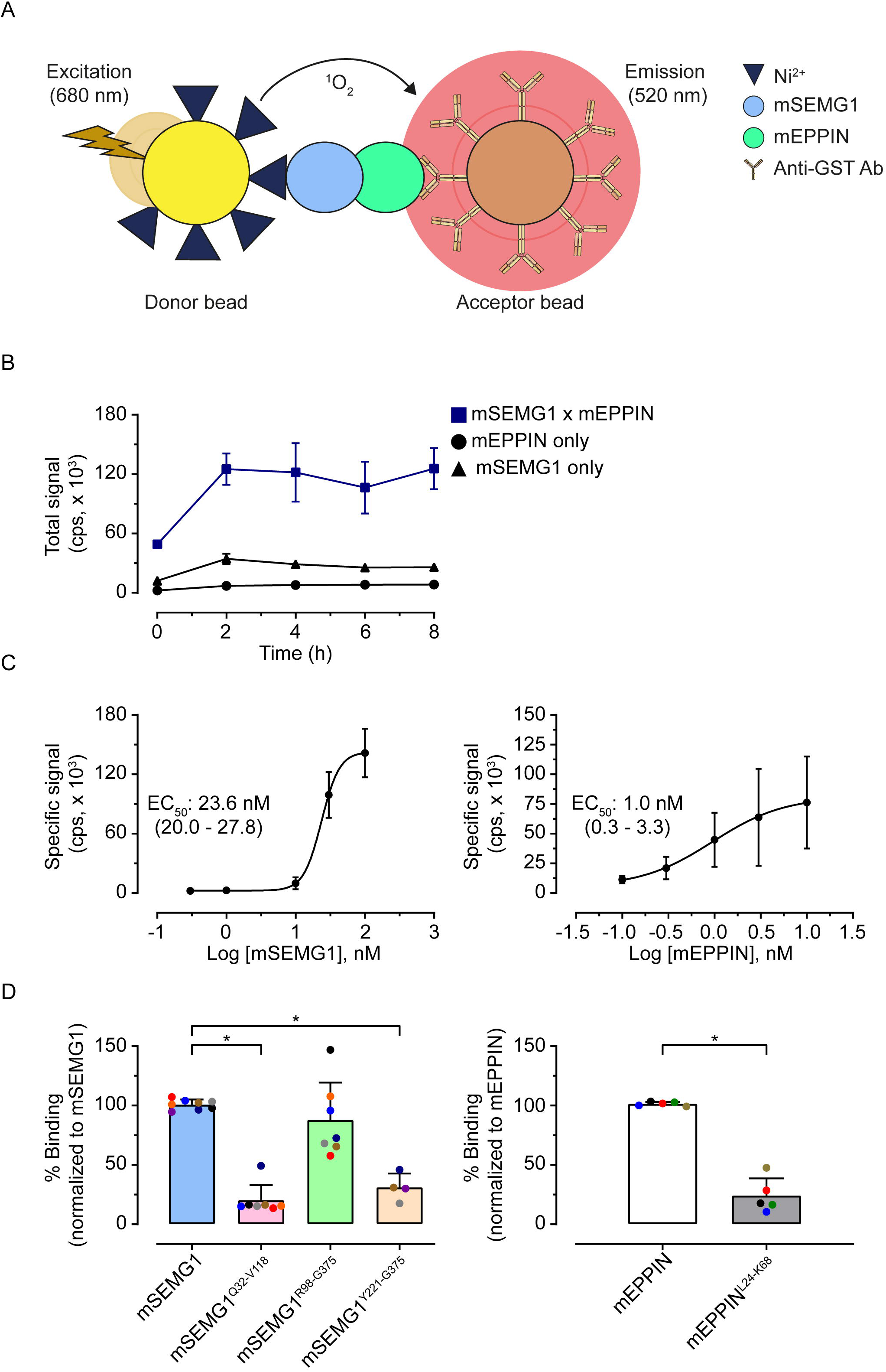
Characterization of the interaction between mSEMG1 and mEPPIN. (A) Schematic representation of the AlphaScreen assay. GST-tagged mEPPIN was captured by anti-GST acceptor beads, and 6×His-tagged mSEMG1 by Ni²⁺-chelate donor beads. Upon excitation at 680 nm, a luminescent signal was generated due to the proximity (<200 nm) of donor and acceptor beads, indicating an interaction between mEPPIN and mSEMG1. (B) mEPPIN was incubated with mSEMG1 in a time-course experiment. Background signal was detected when beads were incubated in the absence of mEPPIN or mSEMG1. Data are presented as mean ± SD from a representative experiment out of ten, each performed in triplicate. (C) Left: Concentration-response curve for mSEMG1 with a constant concentration of mEPPIN (10 nM). Right: Concentration-response curve for mEPPIN with a constant concentration of mSEMG1 (30 nM). A specific signal for each data point was determined by subtracting the background signal from the total signal. Results represent the mean ± SD of specific signals from 2-4 experiments. (D) Left: mSEMG1 and its fragments (Q32-V118: mSEMG1^Q32-V118^, R98-G375: mSEMG1^R98-G375^, and Y221-G375: SEMG^Y221-G375^) were incubated with mEPPIN for 4 h. Right: mEPPIN and its N-terminal construct (D24-K68: mEPPIN^D24-K68^) were incubated with mSEMG1 for 4 h. Dots with the same color represent values from independent experiments. Specific signals for each data point were normalized as a percentage of the signal from full-length mSEMG1/mEPPIN binding. Asterisks indicate statistically significant differences from the control group (mSEMG1 or mEPPIN) (p<0.05, ANOVA followed by Dunnett’s test). Results are expressed as mean ± SD of specific signals from 4-10 experiments. Cps = counts per second.

To map the EPPIN binding site, we tested the mEPPIN binding capacity of our truncated mSEMG1 constructs. mSEMG1^R98-G375^ showed a mEPPIN binding capacity similar to the full-length mSEMG1, both displaying a comparable specific signal (Fig. 6D). In contrast, we observed a reduction in the specific signal by ∼80% (p < 0.0001) and ∼69% (p < 0.0001) for mSEMG1^Q32-V118^ and mSEMG1^Y221-G375^ in comparison to full-length mSEMG1, respectively (Fig. 6D).

In addition, the removal of mouse EPPIN Kunitz domain in a C-terminal truncation lacking residues C69-T134 resulted in a reduction in the specific signal by ∼75% (p = 0.001) compared to that detected with full-length EPPIN (Fig. 6D). Competition assays using anti-EPPIN antibodies Q20E and S21C, that recognized epitopes within EPPIN WFDC and Kunitz domains, respectively, reduced the specific signal of the binding of mEPPIN and mSEMG1 in a concentration-dependent manner with IC_50_ values of 3.0 nM (0.2–34.3 nM) and 95.6 nM (20.6–443.6 nM), respectively (Fig. S5). Altogether, these results demonstrate that the sequence R98-S220 of mSEMG1 contains the major mEPPIN binding domain and that both WFDC and Kunitz domains of mEPPIN are involved in mSEMG1 binding.

## Discussion

During ejaculation, SEMG1 coats the sperm surface by interacting with EPPIN, thereby modulating sperm motility and preventing capacitation-associated events, such as hyperactivation and acrosome exocytosis (O’Rand et al., 2011; Vickram et al., 2022). However, the underlying mechanisms governing the effects of SEMG1 on sperm function remain elusive. Here, we demonstrate that mSEMG1 inhibited sperm hyperactivation at concentrations at least 200-fold lower than those previously reported (Sakaguchi et al., 2020), highlighting SEMG1 as a potent, endogenous inhibitor of sperm hyperactivation. Moreover, our results reveal that mouse SEMG1 contains two distinct functional domains – within the Q32-V118 and R98-S220 sequences – which inhibit both sperm hyperactivation and CatSper currents at similar concentrations. Since only the R98-S220 sequence retained full-extent EPPIN binding capacity relative to full-length mSEMG1, our data indicate that SEMG1 operates through domain-specific mechanisms associated with differential EPPIN interaction.

During capacitation, murine spermatozoa experience a pH_i_ shift from acidic to alkaline, driven by several ion channels and transporters. Among them, Na⁺/H⁺ exchangers (NHEs) are well-established contributors to pHi regulation (Wang et al., 2007; Novero et al., 2024). This event is crucial for the activation of sperm-specific ion channels, such as CatSper, which in turn leads to a rise in [Ca^2+^]_i_ (Lishko and Mannowetz, 2018). Ca^2+^ influx through CatSper plays a fundamental role in reproduction, as channel dysfunction leads to male infertility associated with hyperactivation failure in both humans and mice (Wang et al., 2021). While previous reports have demonstrated that the inhibition of human sperm motility by SEMG1 was related to a decrease in [Ca^2+^]_i_ (O’Rand and Widgren, 2012), whether this effect is associated with CatSper inhibition has not been clarified. As shown herein, mSEMG1 reduces CatSper currents in mouse spermatozoa, an effect not previously reported for this protein, and inhibits hyperactivation at comparable concentrations. Nevertheless, the underlying mechanisms remain to be determined.

Given that intracellular alkalization is a well-established trigger for CatSper activation and sperm hyperactivation (Kirichok et al., 2006), we investigated whether pharmacologically induced pH_i_ increase could rescue hyperactivation following mSEMG1 treatment. Our findings showed that hyperactivated motility remained suppressed despite pH_i_ alkalization, indicating a downstream effect of alkalinization-dependent CatSper activation. During capacitation, increases in [Ca^2+^]_i_ contribute to both Em hyperpolarization and intracellular alkalinization, forming a positive feedback loop that in turn potentiates CatSper activity (Chávez et al., 2014; Novero et al., 2024). In this regard, Em hyperpolarization is mediated by KSper, which localizes along the mouse and human sperm flagellum (Navarro et al., 2007; Lv et al., 2022). mSEMG1 did not produce a detectable change in capacitation-associated Em hyperpolarization at the concentrations tested. Because Em reflects the net contribution of all active conductances, this observation does not establish that KSper activity is preserved; it does, however, bear on the possibility of an mSEMG1-induced acidification, as discussed below.

In the normal physiological context, mouse SEMG1 is present in the seminal plasma at higher concentrations (≥ 25 µM; Kawano and Yoshida, 2007) than those required for inhibiting sperm hyperactivation *in vitro*. Thus, our findings support the model that SEMG1 can prevent premature hyperactivation, consistent with its inhibitory effect on CatSper activity in ejaculated spermatozoa. The mechanism by which mSEMG1 reduces CatSper currents remains to be established, and two non-exclusive possibilities should be considered. mSEMG1 may act indirectly, for instance through intracellular acidification, which would be expected to reduce both CatSper and KSper activity; alternatively, it may act on the channel itself, by occluding cation entry, stabilizing a closed state, or disrupting the organization of the CatSpermasome arrays. Our results do not discriminate between these possibilities, and the reduction in the CatSper currents does not by itself demonstrate a direct interaction between SEMG1 and CatSper.

On the other hand, two observations argue against acidification as the principal mechanism in mouse spermatozoa. First, NH_4_Cl-induced alkalinization did not restore hyperactivated motility in the presence of mSEMG1 (Fig. 2D), placing the inhibitory effect downstream of alkalinization-dependent CatSper activation. Second, murine KSper/SLO3 is exquisitely sensitive to intracellular alkalinization (Navarro et al., 2007; Zeng et al., 2011), unlike its human counterpart, so that mouse sperm Em is a sensitive, if indirect, reporter of pHi. Thus, a substantial mSEMG1-induced acidification would be expected to depolarize capacitated spermatozoa, which we did not observe. This inference remains indirect, as pHi was not measured in mouse spermatozoa, and SEMG1-dependent pHi changes have so far been reported only in human spermatozoa (O’Rand and Widgren, 2012). A further limitation concerns the Em measurements. Em was determined by population fluorimetry with DiSC₃(5), using an internal valinomycin/K⁺ calibration curve generated for each individual determination, an approach previously validated in mammalian spermatozoa (Ritagliati et al., 2018; Baro Graf et al., 2020). This method reports a population average and lacks single-cell resolution, and is therefore less sensitive than current-clamp recordings to small or heterogeneous voltage changes. Direct current-clamp recordings, together with direct measurement of pHi in mouse spermatozoa exposed to mSEMG1, will be required to define the mechanism of CatSper inhibition.

Additionally, it is possible that higher mSEMG1 concentrations are required to modulate sperm Em. Mouse SEMG1 undergoes proteolytic cleavage in the female reproductive tract (Kawano and Yoshida, 2007); thus, it is plausible that different concentrations of mSEMG1 are required to modulate distinct events associated with sperm capacitation. Another putative mechanism involves the ability of SEMG1 to stabilize membrane microdomains through modulation of cholesterol removal and binding to lipid raft components, such as the ganglioside GM1 (Kawano et al., 2008). Further experiments, including evaluation of sperm intracellular pH and KSper activity, are required to establish the effects of mSEMG1 on these parameters.

Our findings identified two functional domains within mSEMG1, spanning Q32-V118 and R98-S220 sequences, that play crucial roles in inhibiting sperm hyperactivation. Truncated constructs containing these sequences promoted inhibition of CatSper activity to a similar extent as the full-length mSEMG1. These findings are consistent with previous observations that SEMG1 is a modular protein, containing discrete domains that contribute to its multifunctional roles in sperm function and protection (de Lamirande, 2007; Vickram et al., 2022). Upon ejaculation, SEMG1 associates with the sperm surface through binding to EPPIN, a process known to be critical for regulating sperm motility (Mitra et al., 2010; O’Rand et al., 2011; O’Rand and Widgren, 2012). Our previous results demonstrated the co-immunoprecipitation of EPPIN and SEMG1 from mouse spermatozoa incubated with the seminal vesicle fluid when either anti-EPPIN or anti-SEMG1 antibodies were used, indicating a parallel system to humans (Wang et al., 2005; Mariani et al., 2020). These observations prompted us to investigate the relevance of EPPIN binding to the effects of mSEMG1 on sperm function. Our data showed that mouse EPPIN and SEMG1 are binding partners, confirming that EPPIN is a docking site for SEMG1 on the mouse sperm surface, and further demonstrated functional conservation of this interaction by demonstrating that mouse EPPIN can also bind human SEMG1.

We identified the mSEMG1 R98-S220 sequence, which lacks its unique cysteine residue but encompasses two long amino acid repeats, as containing the major EPPIN binding domain. This result was unexpected, since human SEMG1 relies on amino acid sequences flanking its unique cysteine residue to bind EPPIN and inhibit sperm motility (Silva et al., 2013). This divergence may reflect the rapid adaptive evolution of *SEMG1* genes across species (Ferreira et al., 2013; Silva et al., 2013), which could shift EPPIN-binding pockets to different regions of the primary sequence when mouse and human SEMG1 are compared. Conversely, the mSEMG1 Q32-V118 sequence, which contains its unique cysteine residue, displayed low EPPIN binding *in vitro* (<20% relative to the full-length mSEMG1), yet retained the ability to inhibit CatSper currents and sperm hyperactivation. These findings indicate that different mSEMG1 domains inhibit sperm hyperactivation and that strong EPPIN binding is not strictly required for CatSper inhibition by this SEMG1 fragment once present at the sperm surface. Nevertheless, we cannot rule out the possibility that EPPIN binding may facilitate the initial positioning of both mSEMG1 R98-S220 and Q32-V118 sequences on the sperm surface, enabling distinct SEMG1 domains to engage downstream targets, such as CatSper. Importantly, recent genetic studies show that the combined ablation of *Eppin* and its paralogs *Wfdc6a*, *Wfdc8*, and *Wfcd6b* in mice led to an azoospermic phenotype, underscoring the essential role of EPPIN in sperm development but precluding direct functional testing of mSEMG1 effects in mature spermatozoa lacking EPPIN (Kent et al., 2024). Altogether, we conclude that mouse SEMG1 may inhibit CatSper and hyperactivation through domain-specific mechanisms associated with differential EPPIN-binding capacity.

Based on our findings, we propose a working model by which SEMG1 regulates hyperactivation in ejaculated spermatozoa in mice (Fig. 7). Following ejaculation, SEMG1 coats the surface of spermatozoa through EPPIN binding in the semen coagulum/copulatory plug, preventing premature hyperactivation and capacitation within the female reproductive tract. This event is mediated by two distinct SEMG1 functional domains within Q32-V118 and R98-S220 sequences that inhibit CatSper activity through domain-specific mechanisms. In the oviduct, a putative PSA-like protease (likely belonging to the kallikrein family) cleaves SEMG1, allowing spermatozoa to undergo hyperactivation and capacitation. Collectively, these findings position SEMG1 Q32-V118 and R98-S220 sequences as compelling prototypes for the rational design of CatSper inhibitors with potential applications in male contraception targeting critical regulatory pathways for hyperactivation.

**Figure 7.**
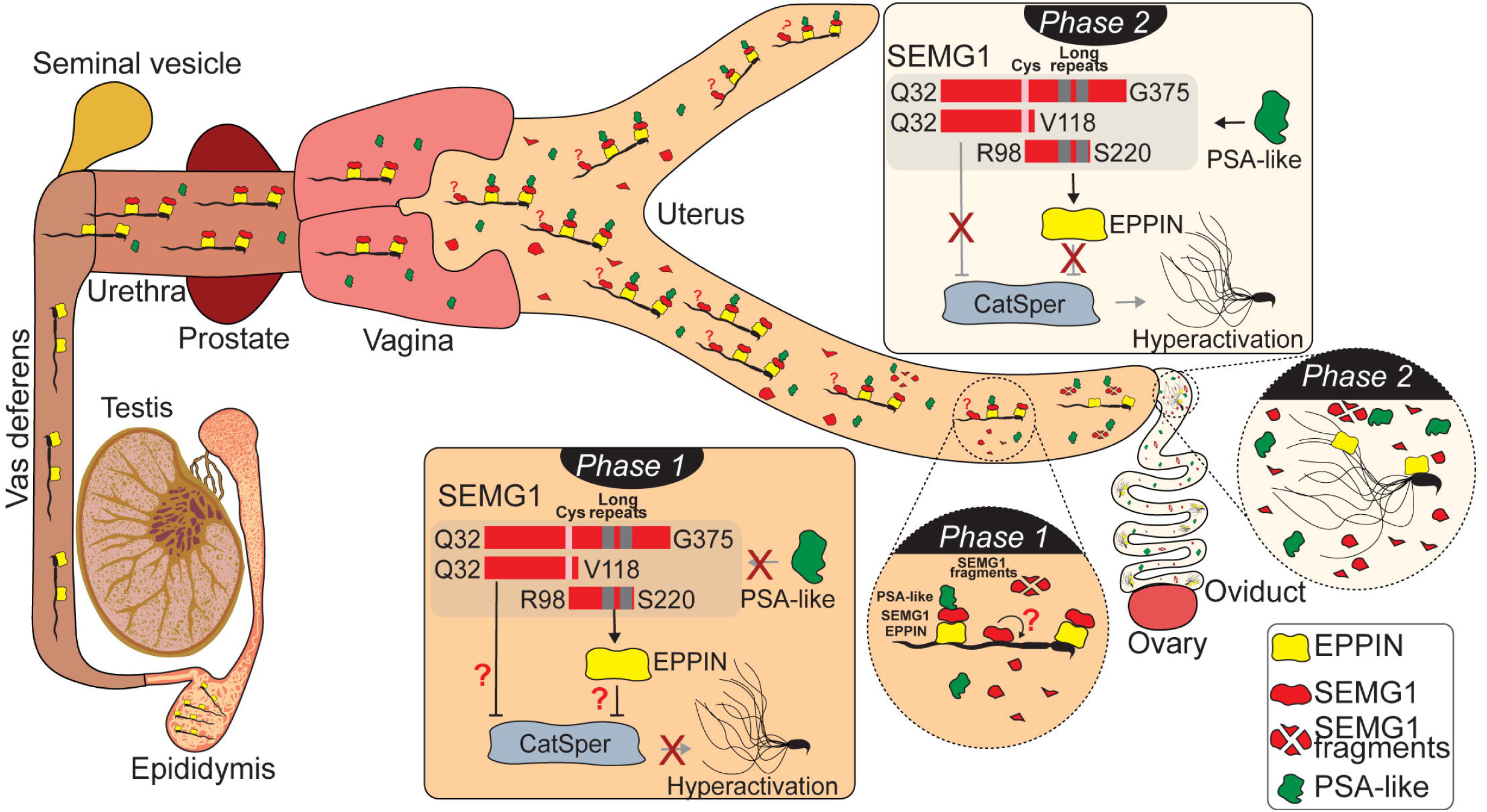
Working model of SEMG1-mediated inhibition of sperm hyperactivation after sperm ejaculation. EPPIN is present on the surface of mature spermatozoa stored in the cauda epididymidis. In the semen coagulum/copulatory plug (Phase 1), SEMG1 binds to the sperm surface via interaction with EPPIN through a binding domain within residues R98-S220, thus coating spermatozoa. SEMG1 may also interact with spermatozoa independently of EPPIN binding. Domains within the SEMG1 Q32-G375 and R98-S220 regions inhibit sperm function, i.e., temporarily suppressing hyperactivation and CatSper currents in the uterus, while only the R98-S220 domain contains the EPPIN binding site. As spermatozoa move to the upper regions of the female reproductive tract, SEMG1 is proteolytically cleaved by a PSA-like enzyme (Phase 2), whose identity remains unknown in mice, allowing hyperactivation to take place in the oviduct. Question marks indicate mechanisms that remain to be elucidated.

### Experimental Procedures

#### Animals

We used male (90–120 days old) Swiss mice obtained from the Center for Research and Production of Laboratory Animals (CPPA) of São Paulo State University, Botucatu, SP, Brazil. The animals were maintained under controlled conditions (12□h light/dark cycle, 22–24°C) with food and water available ad libitum. All procedures were approved by the Animal Care and Use Committee of UNESP (protocol #5219150420) and followed the guidelines of the Brazilian National Council for the Control of Animal Experimentation (CONCEA). We also used wild-type (WT) and CatSper1-knockout (KO) male C57BL/6 mice, maintained at the Animal Facility of Washington University in St. Louis, School of Medicine, St. Louis, MO, United States (protocol #22-0251), and the Facultad de Ciencias Bioquímicas y Farmacéuticas de Rosario, Rosario, SF, Argentina (protocol #492/2018). These animals were used for patch-clamp recordings, membrane potential analysis, and additional sperm motility experiments. All experiments were conducted following the institutional guidelines for animal research and were approved by the respective Animal Care and Use Committees. Mice were euthanized by inhaled isoflurane overdose or CO_2_ inhalation, followed by cervical dislocation.

#### Recombinant proteins

Recombinant full-length 6xHis-mSEMG1 (Q32-G375), its truncated fragments 6×His-mSEMG1^Q32–V118^ (Q32–V118); 6×His-mSEMG1^R98–G375^ (R98–G375); 6×His-mSEMG1^Y221–G375^ (Y221–G375), as well as full-length mEPPIN (S23-T134) and its N-terminal fragment mEPPIN^D24-K68^ (D24-K68) were purchased from GenScript (Piscataway, NJ, USA). Proteins were expressed in E. coli using the pET-30a(+) expression system and purified by sequential Ni-affinity and Superdex 200 chromatography. Protein concentrations were determined by Bradford assay using BSA as a standard. Purified proteins were supplied in PBS containing 10% glycerol and 0.5 M L-arginine (pH 7.4) and stored at −80°C until use. According to the manufacturer’s quality-control analysis, protein purity was above 90% as determined by densitometric analysis of Coomassie Blue-stained SDS-PAGE gels. In addition, protein identity and purity were independently validated in our laboratory by Amido Black staining and Western blot (Fig. S6). Before each experimental procedure, the recombinant proteins were dialyzed using 3.5 MWCO mini dialysis cassettes (ThermoFisher Scientific, cat. #69580) against the corresponding experimental buffer: DVF bath solution for electrophysiological recordings and HTF medium for sperm motility analyses. This dialysis step was performed to minimize the carryover of components from the protein storage solution, particularly PBS and other potentially interfering constituents, into the experimental preparations.

#### Protein-protein interaction assays

We performed AlphaScreen to evaluate protein–protein interactions as previously described (Silva et al., 2012), with the following modifications. Briefly, recombinant mEPPIN and its N-terminal fragment, both containing an N-terminal GST-tag, were pre-incubated with anti-GST acceptor beads for 60 minutes. In parallel, mSEMG1 and its truncated fragments (Fig. 4A), containing a 6x-His tag, were incubated with Ni-NTA-chelate donor beads under the same conditions. Equal volumes of the mEPPIN/acceptor bead suspension and the mSEMG1/donor bead suspension were transferred to white opaque 384-well microplates (OptiPlate-384; PerkinElmer, Waltham, MA), in a final reaction volume of 30□µL per well. Plates were sealed and placed in a Synergy 2 Multiplatform automated plate reader (BioTek, Winooski, VT). After 2 min of shaking, readings were taken every 2 h over 8 h, using a 680/30 excitation filter and a 570/100 emission filter. Data acquisition was performed using Gen5 software (BioTek), and a total of 9 time points were recorded per experiment. All conditions were assayed in triplicate.

Final concentrations of the assay components ranged from 0.1-10□nM for mEPPIN, 0.3-100□nM for mSEMG1, and 10□µg/mL for the donor and acceptor beads. Negative controls were conducted under the same conditions in the absence of either mEPPIN or mSEMG1, with beads only. The specific signal at each time point was calculated by subtracting the background signal (obtained in the absence of one of the proteins) from the total signal. To determine the EC_50_ for mEPPIN binding to mSEMG1, we applied nonlinear regression curve fitting to the specific signal values obtained after 4 h of incubation. For competition assays, we incubated mEPPIN (10□nM) and mSEMG1 (30□nM) in the presence of increasing concentrations of anti-EPPIN Q20E (targeting the P20–E39 epitope) or anti-EPPIN S21C (targeting the S103–C121 epitope) antibodies, maintaining a total reaction volume of 30□µL and bead concentration of 10□µg/mL. Specific signals for each competitor concentration were calculated as described, and IC_50_ values were determined by nonlinear regression.

#### Analysis of Sperm Motility

Analysis of sperm motility was performed as previously described (Silva et al., 2021), with the following modifications. We collected mouse spermatozoa from the cauda epididymides and incubated them in human tubal fluid (HTF) medium at 37□°C under 5% CO_2_. After 10 min of incubation, we removed the tissue and diluted the sperm suspension (∼40,000 cells) in HTF supplemented with 25□mM sodium bicarbonate and 0.75% (m/v) bovine serum albumin (BSA; Sigma, cat. #SLBT1197). We treated the sperm suspensions with recombinant mSEMG1 or its truncated constructs (1-25□μM), with the highest concentration corresponding to the physiological range previously reported for SEMG1 (Kawano and Yoshida, 2007) (Fig. 4). Negative controls were prepared using HTF medium without recombinant proteins. Rescue experiments were performed to assess whether intracellular alkalinization could counteract the inhibitory effects of mSEMG1 on sperm motility. Alkalinization was induced by adding NH_4_Cl, and its efficacy was verified by analyzing sperm motility under non-capacitating conditions (medium lacking bicarbonate and BSA) after NH_4_Cl treatment. To evaluate the effects of mSEMG1 and its truncated constructs on sperm motility, the recombinant proteins were dialyzed against the HTF medium before use to minimize the carryover of the protein storage buffer into the analysis. Sperm motility was analyzed using a computer-assisted sperm analysis (CASA) system (Hamilton-Thorne, Beverly, MA, USA) with Animal Motility II Software version 1.9. We acquired images using an Olympus CX41 microscope equipped with a MiniTherm stage warmer and a JAI CM-040 digital camera. Aliquots of the sperm suspensions were loaded into pre-warmed 80-μm 2X-CEL slides (Hamilton-Thorne) at 37°C using large-bore pipette tips. We assessed sperm motility at 20-120 min after incubation.

We recorded sperm tracks for 1.5□s (90 frames) at a frame rate of 60□Hz using a 40× negative-phase contrast objective. For each sample, we analyzed at least 200 spermatozoa across 10 different fields. We used video playback to review and manually correct the tracks, removing mistracked cells caused by collisions or debris interference. We then extracted individual ASCII files containing the kinematic parameters for each sperm cell. The following parameters were analyzed: average path velocity (VAP, µm/s), straight-line velocity (VSL, µm/s), curvilinear velocity (VCL, µm/s), amplitude of lateral head displacement (ALH, μm), straightness (STR, %), and linearity (LIN, %). We classified sperm as progressive when VAP□>□50 µm s^-1^ and STR□>□50%, hyperactivated when□VCL >□180□µm s^-1^, ALH >□9.5□μm, and LIN <□23.6%, and slow when VAP ≤ 30□µm s^-1^ and VSL ≤□50□µm s^-1^ (Silva et al., 2021). To assess the potential effects of mSEMG1 and its fragments on sperm viability after the maximum incubation period of 120 min under capacitating conditions, viability was evaluated in parallel with motility assays using eosin-nigrosin staining. The hyperactivation inhibition rate was determined from the percentage of hyperactivated spermatozoa normalized to its respective control group within each independent experiment, calculated as (% hyperactivated spermatozoa in the treated sample divided by % hyperactivated spermatozoa in the matched control) × 100. The resulting values were fitted by non-linear regression (best-fit) to obtain the corresponding IC_50_. The progressive motility inhibition rate was determined using the same approach.

#### Electrophysiology

Electrophysiological recordings were performed as previously described (Liu et al., 2021). In summary, gigaohm seals were established at the cytoplasmic droplet of mouse sperm (Fig. 3) using HS containing (in mM): 130 NaCl, 5 KCl, 1 MgSO_4_, 2 CaCl_2_, 5 glucose, 1 sodium pyruvate, 10 lactic acid, and 20 HEPES (pH 7.4). Whole-cell configuration was achieved by applying brief (1 ms) voltage pulses of 400–630 mV combined with gentle suction. Currents were elicited with voltage ramps from −80□mV to +80□mV, starting from a holding potential of 0□mV, in both HS and DVF bath solutions. Data were acquired at a sampling rate of 2–5□kHz using a low-pass filter at 1□kHz. Patch pipettes (11–17□MΩ) were filled with an internal solution containing (in mM): 130 Cs-methanesulfonate, 70 HEPES, 3 EGTA, 2 EDTA, and 0.5 Tris-HCl (pH adjusted to 7.4 with CsOH). The DVF bath solution used for recording monovalent CatSper currents contained (in mM): 140 Cs-methanesulfonate, 40 HEPES, and 1 EDTA (pH 7.4, adjusted with CsOH). Leak currents were recorded in HS solution and remained close to zero. These currents were subtracted from the currents recorded in DVF, both in the absence and presence of the proteins, before data analysis. To evaluate the effects of mSEMG1 and its truncated constructs on I*_CatSper_*, the recombinant proteins were dialyzed against the DVF bath solution before use to minimize the carryover of the protein storage buffer into the electrophysiological recordings. To confirm the efficiency of this procedure, I*_CatSper_* was recorded after dialysis of the protein storage buffer against DVF. The similar current traces recorded in DVF and dialyzed storage buffer indicate that the dialysis procedure did not alter I*_CatSper_* (Fig. S7). During recordings, bath perfusion was temporarily stopped, and the chamber volume was maintained at approximately 400–500 μL. Proteins were then added directly to the recording chamber to achieve final concentrations of 5 or 10 μM. The complete compositions of the HS, DVF, and pipette solutions are listed in Tables S1, S2, and S3. All electrophysiology recordings were conducted at Washington University in St. Louis, School of Medicine, St. Louis, MO, USA.

#### Membrane Potential assays

Changes in sperm Em were assessed using DiSC3(5) (Molecular Probes, cat. #D306), as previously described (Ritagliati et al., 2018). A sperm suspension aliquot (∼4.0 x 10□cells) was diluted in TYH medium supplemented with 20 mM sodium bicarbonate and 0.5% (m/v) BSA, with or without mSEMG1 and its fragments (10 μM). Capacitated and non-capacitated sperm in the absence of mSEMG1 were used as experimental controls. After treatment with the proteins and 30 min of capacitation, cells were stained with the Em-sensitive dye DiSC3(5) 1 μM (Molecular Probes) for 2 min. Spermatozoa were transferred to a cuvette at 37°C with mild agitation, and fluorescence was monitored using a Cary Eclipse fluorescence spectrophotometer (Agilent, CA) at excitation/emission wavelengths of 620/670 nm. Recordings began when steady-state fluorescence was reached, and calibration was performed at the end of each measurement by adding 1 μM valinomycin with sequential KCl additions for internal calibration curves. Sperm Em was derived from initial fluorescence (measured in arbitrary fluorescence units) by linear interpolation of theoretical Em values from the calibration curve against arbitrary fluorescence units for each trace. This internal calibration for each determination compensated for variables influencing absolute fluorescence values. To evaluate the effects of mSEMG1 on Em, the recombinant protein was dialyzed against the TYH medium before use to minimize the carryover of the protein storage buffer into the recordings.

### Statistical Analysis

Data are presented as mean ± standard deviation (SD). Normality and homoscedasticity were assessed using the Kolmogorov–Smirnov test and F-test, respectively. Statistical differences between two independent groups were analyzed using an unpaired Student’s t-test; for comparisons involving more than two groups, one-way ANOVA followed by Dunnett’s post hoc test was used. For electrophysiological recordings, a paired Student’s t-test was applied. Percentage data were arcsine square root transformed before statistical analysis (Zar, 2010). A p-value < 0.05 was considered statistically significant.

## Acknowledgments

We thank Luiz Oliveira, Paulo Mioni, and Flávia Colenci, Ph.D., Department of Biophysics and Pharmacology, IBB/UNESP, for their technical assistance. We are also grateful to Michael ÓRand, Ph.D., and Katherine Hamil, M.Sc., for their critical reading of the manuscript and valuable insights, and Célio Fernandes, Ph.D., for his helpful discussions during the project.

## Data and resource availability

All relevant data and details of resources can be found within the article and its supplementary information. Any material (e.g., plasmids and recombinant proteins) generated in this paper can be obtained by emailing the corresponding author.

## References

Araki, N., Trencsényi, G., Krasznai, Z. T., Nizsalóczki, E., Sakamoto, A., Kawano, N., Miyado, K., Yoshida, K. and Yoshida, M. (2015). Seminal Vesicle Secretion 2 acts as a protectant of sperm sterols and prevents ectopic sperm capacitation in mice. Biol. Reprod. 92(1), 8, 1-10-18, 11-10. doi:10.1095/biolreprod.114.120642

Austin, C. R. (1952). The ‘Capacitation’ of the mammalian sperm. Nature 170(4321), 326–326. doi:10.1038/170326a0

Avenarius, M. R., Hildebrand, M. S., Zhang, Y., Meyer, N. C., Smith, L. L., Kahrizi, K., Najmabadi, H. and Smith, R. J. (2009). Human male infertility caused by mutations in the CATSPER1 channel protein. Am. J. Hum. Genet. 84(4), 505–510. doi:10.1016/j.ajhg.2009.03.004

Baro Graf, C., Ritagliati, C., Torres-Monserrat, V., Stival, C., Carizza, C., Buffone, M. G. and Krapf, D. (2020). Membrane Potential Assessment by Fluorimetry as a Predictor Tool of Human Sperm Fertilizing Capacity. Frontiers in Cell and Developmental Biology 7(383). doi:10.3389/fcell.2019.00383

Bystroff, C. (2018). Intramembranal disulfide cross-linking elucidates the super-quaternary structure of mammalian CatSpers. Reprod. Biol. 18(1), 76–82. doi:10.1016/j.repbio.2018.01.005

Cancel, A. M., Lobdell, D., Mendola, P. and Perreault, S. D. (2000). Objective evaluation of hyperactivated motility in rat spermatozoa using computer-assisted sperm analysis. Human. Reprod. 15(6), 1322–1328. doi:10.1093/humrep/15.6.1322

Carlson, A. E., Quill, T. A., Westenbroek, R. E., Schuh, S. M., Hille, B. and Babcock, D. F. (2005). Identical phenotypes of CatSper1 and CatSper2 null sperm. J. Biol. Chem. 280(37), 32238–32244. doi:10.1074/jbc.M501430200

Chang, M. C. (1951). Fertilizing capacity of spermatozoa deposited into the fallopian tubes. Nature 168, 697. doi:10.1038/168697b0

Chávez, J. C., Ferreira, J. J., Butler, A., De La Vega Beltrán, J. L., Treviño, C. L., Darszon, A., Salkoff, L. and Santi, C. M. (2014). SLO3 K^+^ channels control calcium entry through CATSPER channels in sperm. J. Biol. Chem. 289(46), 32266–32275. doi:10.1074/jbc.M114.607556

Chung, J.-J., Miki, K., Kim, D., Shim, S.-H., Shi, H. F., Hwang, J. Y., Cai, X., Iseri, Y., Zhuang, X. and Clapham, D. E. (2017). CatSperζ regulates the structural continuity of sperm Ca^2+^ signaling domains and is required for normal fertility. eLife 6, e23082. doi:10.7554/eLife.23082

Darszon, A., Nishigaki, T., Beltran, C. and Treviño, C. L. (2011). Calcium channels in the development, maturation, and function of spermatozoa. Physiol. Rev. 91(4), 1305–1355. doi:10.1152/physrev.00028.2010

De Jonge, C. (2017). Biological basis for human capacitation-revisited. *Human Reprod*. Update 23(3), 289–299. doi:10.1093/humupd/dmw048

de Lamirande, E. (2007). Semenogelin, the main protein of the human semen coagulum, regulates sperm function. Semin Thromb Hemost 33(1), 60–68. doi:10.1055/s-2006-958463

de Lamirande, E. and Lamothe, G. (2010). Levels of semenogelin in human spermatozoa decrease during capacitation: involvement of reactive oxygen species and zinc. Human. Reprod. 25(7), 1619–1630. doi:10.1093/humrep/deq110

Ferreira, Z., Seixas, S., Andrés, A. M., Kretzschmar, W. W., Mullikin, J. C., Cherukuri, P. F., Cruz, P., Swanson, W. J., Program, N. C. S., Clark, A. G., Green, E. D. and Hurle, B. (2013). Reproduction and Immunity-Driven Natural Selection in the Human WFDC Locus. Molecular Biology and Evolution 30(4), 938–950. doi:10.1093/molbev/mss329

Freitas, M. J., Vijayaraghavan, S. and Fardilha, M. (2017). Signaling mechanisms in mammalian sperm motility. Biol. Reprod. 96(1), 2–12. doi:10.1095/biolreprod.116.144337

Hildebrand, M. S., Avenarius, M. R., Fellous, M., Zhang, Y., Meyer, N. C., Auer, J., Serres, C., Kahrizi, K., Najmabadi, H., Beckmann, J. S. and Smith, R. J. (2010). Genetic male infertility and mutation of CATSPER ion channels. Eur. J. Hum. Genet. 18(11), 1178–1184. doi:10.1038/ejhg.2010.108

Hwang, J. Y., Mannowetz, N., Zhang, Y., Everley, R. A., Gygi, S. P., Bewersdorf, J., Lishko, P. V. and Chung, J.-J. (2019). Dual sensing of physiologic pH and calcium by EFCAB9 regulates sperm motility. Cell 177(6), 1480–1494.e1419. doi:10.1016/j.cell.2019.03.047

Katz, D. F. and Yanagimachi, R. (1980). Movement characteristics of hamster spermatozoa within the oviduct. Biol. Reprod. 22(4), 759–764. doi:10.1095/biolreprod22.4.759

Kawano, N., Araki, N., Yoshida, K., Hibino, T., Ohnami, N., Makino, M., Kanai, S., Hasuwa, H., Yoshida, M., Miyado, K. and Umezawa, A. (2014). Seminal vesicle protein SVS2 is required for sperm survival in the uterus. Proc. Natl. Acad. Sci. USA 111(11), 4145–4150. doi:10.1073/pnas.1320715111

Kawano, N., Yoshida, K., Iwamoto, T. and Yoshida, M. (2008). Ganglioside GM1 mediates decapacitation effects of SVS2 on murine spermatozoa. Biol. Reprod. 79(6), 1153–1159. doi:10.1095/biolreprod.108.069054

Kawano, N. and Yoshida, M. (2007). Semen-coagulating protein, SVS2, in mouse seminal plasma controls sperm fertility. Biol. Reprod. 76(3), 353–361. doi:10.1095/biolreprod.106.056887

Kent, K., Nozawa, K., Parkes, R., Dean, L., Daniel, F., Leng, M., Jain, A., Malovannaya, A., Matzuk, M. M. and T, X. G. (2024). Large-scale CRISPR/Cas9 deletions within the WFDC gene cluster uncover gene functionality and critical roles in mammalian reproduction. Proc. Natl. Acad. Sci. USA 121(51), e2413195121. doi:10.1073/pnas.2413195121

Kirichok, Y. and Lishko, P. V. (2011). Rediscovering sperm ion channels with the patch-clamp technique. Mol. Hum. Reprod. 17(8), 478–499. doi:10.1093/molehr/gar044

Kirichok, Y., Navarro, B. and Clapham, D. E. (2006). Whole-cell patch-clamp measurements of spermatozoa reveal an alkaline-activated Ca^2+^ channel. Nature 439(7077), 737–740. doi:10.1038/nature04417

Lin, S., Ke, M., Zhang, Y., Yan, Z. and Wu, J. (2021). Structure of a mammalian sperm cation channel complex. Nature 595(7869), 746–750. doi:10.1038/s41586-021-03742-6

Lishko, P. V. and Mannowetz, N. (2018). CatSper: a unique calcium channel of the sperm flagellum. Curr. Opin. Physiol. 2, 109–113. doi:10.1016/j.cophys.2018.02.004

Liu, B., Mundt, N., Miller, M., Clapham, D. E., Kirichok, Y. and Lishko, P. V. (2021). Recording electrical currents across the plasma membrane of mammalian sperm cells. J. Vis. Exp. 168. doi:10.3791/62049

López-González, I., Torres-Rodríguez, P., Sánchez-Carranza, O., Solís-López, A., Santi, C. M., Darszon, A. and Treviño, C. L. (2014). Membrane hyperpolarization during human sperm capacitation. Mol. Hum. Reprod. 20(7), 619–629. doi:10.1093/molehr/gau029

Lv, M., Liu, C., Ma, C., Yu, H., Shao, Z., Gao, Y., Liu, Y., Wu, H., Tang, D., Tan, Q., Zhang, J., Li, K., Xu, C., Geng, H., Zhang, J., Li, H., Mao, X., Ge, L., Fu, F., Zhong, K., Xu, Y., Tao, F., Zhou, P., Wei, Z., He, X., Zhang, F. and Cao, Y. (2022). Homozygous mutation in SLO3 leads to severe asthenoteratozoospermia due to acrosome hypoplasia and mitochondrial sheath malformations. Reprod Biol Endocrinol 20(1), 5. doi:10.1186/s12958-021-00880-4

Mariani, N. A. P., Camara, A. C., Silva, A. A. S., Raimundo, T. R. F., Andrade, J. J., Andrade, A. D., Rossini, B. C., Marino, C. L., Kushima, H., Santos, L. D. and Silva, E. J. R. (2020). Epididymal protease inhibitor (EPPIN) is a protein hub for seminal vesicle-secreted protein SVS2 binding in mouse spermatozoa. Mol. Cell. Endocrinol. 506, 110754. doi:10.1016/j.mce.2020.110754

Mitra, A., Richardson, R. T. and O’Rand, M. G. (2010). Analysis of recombinant human Semenogelin as an inhibitor of human sperm motility. Biol. Reprod. 82(3), 489–496. doi:10.1095/biolreprod.109.081331

Navarro, B., Kirichok, Y. and Clapham, D. E. (2007). KSper, a pH-sensitive K^+^ current that controls sperm membrane potential. Proc. Natl. Acad. Sci. USA 104(18), 7688–7692. doi:10.1073/pnas.0702018104

Novero, A. G., Torres Rodríguez, P., De la Vega Beltrán, J. L., Schiavi-Ehrenhaus, L. J., Luque, G. M., Carruba, M., Stival, C., Gentile, I., Ritagliati, C., Santi, C. M., Nishigaki, T., Krapf, D., Buffone, M. G., Darszon, A., Treviño, C. L. and Krapf, D. (2024). The sodium–proton exchangers sNHE and NHE1 control plasma membrane hyperpolarization in mouse sperm. J. Biol. Chem. 300(12), 107932. doi:10.1016/j.jbc.2024.107932

O’Rand, M. G. and Widgren, E. E. (2012). Loss of calcium in human spermatozoa via EPPIN, the semenogelin receptor. Biol. Reprod. 86(2), 55, 51-57. doi:10.1095/biolreprod.111.094227

O’Rand, M. G., Widgren, E. E., Hamil, K. G., Silva, E. J. and Richardson, R. T. (2011). Epididymal protein targets: A brief history of the development of epididymal protease inhibitor as a contraceptive. J. Androl. 32(6), 698–704. doi:10.2164/jandrol.110.012781

Quill, T. A., Sugden, S. A., Rossi, K. L., Doolittle, L. K., Hammer, R. E. and Garbers, D. L. (2003). Hyperactivated sperm motility driven by CatSper2 is required for fertilization. Proc. Natl. Acad. Sci. USA 100(25), 14869–14874. doi:10.1073/pnas.2136654100

Ren, D., Navarro, B., Perez, G., Jackson, A. C., Hsu, S., Shi, Q., Tilly, J. L. and Clapham, D. E. (2001). A sperm ion channel required for sperm motility and male fertility. Nature 413(6856), 603–609.

Ritagliati, C., Baro Graf, C., Stival, C. and Krapf, D. (2018). Regulation mechanisms and implications of sperm membrane hyperpolarization. Mech. Dev. 154, 33–43. doi:10.1016/j.mod.2018.04.004

Ritagliati, C., Luque, G. M., Stival, C., Baro Graf, C., Buffone, M. G. and Krapf, D. (2018). Lysine acetylation modulates mouse sperm capacitation. Scientific Reports 8(1), 13334. doi:10.1038/s41598-018-31557-5

Robert, M. and Gagnon, C. (1999). Semenogelin I: a coagulum forming, multifunctional seminal vesicle protein. Cell. Mol. Life Sci. 55(6), 944–960.

Sakaguchi, D., Miyado, K., Iwamoto, T., Okada, H., Yoshida, K., Kang, W., Suzuki, M., Yoshida, M. and Kawano, N. (2020). Human Semenogelin 1 promotes sperm survival in the mouse female reproductive tract. Int. J. Mol. Sci. 21(11). doi:10.3390/ijms21113961

Schreiber, M., Wei, A., Yuan, A., Gaut, J., Saito, M. and Salkoff, L. (1998). Slo3, a novel pH-sensitive K+ channel from mammalian spermatocytes. J Biol Chem 273(6), 3509–3516. doi:10.1074/jbc.273.6.3509

Silva, A. A. S., Raimundo, T. R. F., Mariani, N. A. P., Kushima, H., Avellar, M. C. W., Buffone, M. G., Paula-Lopes, F. F., Moura, M. T. and Silva, E. J. R. (2021). Dissecting EPPIN protease inhibitor domains in sperm motility and fertilizing ability: repercussions for male contraceptive development. Mol. Hum. Reprod. 27(12), 1–15. doi:10.1093/molehr/gaab066

Silva, E. J. R., Hamil, K. G. and O’Rand, M. G. (2013). Interacting proteins on human spermatozoa: Adaptive evolution of the binding of semenogelin I to EPPIN. PLoS One 8(12), e82014. doi:10.1371/journal.pone.0082014

Silva, E. J. R., Hamil, K. G., Richardson, R. T. and O’Rand, M. G. (2012). Characterization of EPPIN’s Semenogelin I binding site: A contraceptive drug target. Biol. Reprod. 87(3), 56, 51-58. doi:10.1095/biolreprod.112.101832

Stival, C., Puga Molina, L. d. C., Paudel, B., Buffone, M. G., Visconti, P. E. and Krapf, D. (2016). Sperm capacitation and acrosome reaction in mammalian sperm. Sperm Acrosome Biogenesis and Function During Fertilization (ed. Buffone, M. G.), pp. 93–106. Switzerland: Springer Cham.

Suárez, S. S. and Osman, R. A. (1987). Initiation of hyperactivated flagellar bending in mouse sperm within the female reproductive tract. Biol. Reprod. 36(5), 1191–1198. doi:10.1095/biolreprod36.5.1191

Vickram, S., Rohini, K., Anbarasu, K., Dey, N., Jeyanthi, P., Thanigaivel, S., Issac, P. K. and Arockiaraj, J. (2022). Semenogelin, a coagulum macromolecule monitoring factor involved in the first step of fertilization: A prospective review. Int. J. Biol. Macromol. 209, 951–962. doi:10.1016/j.ijbiomac.2022.04.079

Vyklicka, L. and Lishko, P. V. (2020). Dissecting the signaling pathways involved in the function of sperm flagellum. Curr. Opin. Cell Biol. 63, 154–161. doi:10.1016/j.ceb.2020.01.015

Wang, D., Hu, J., Bobulescu, I. A., Quill, T. A., McLeroy, P., Moe, O. W. and Garbers, D. L. (2007). A sperm-specific Na^+^/H^+^ exchanger (sNHE) is critical for expression and *in vivo* bicarbonate regulation of the soluble adenylyl cyclase (sAC). Proc. Natl. Acad. Sci. USA 104(22), 9325–9330. doi:10.1073/pnas.0611296104

Wang, H., McGoldrick, L. L. and Chung, J.-J. (2021). Sperm ion channels and transporters in male fertility and infertility. Nat. Rev. Urol. 18(1), 46–66. doi:10.1038/s41585-020-00390-9

Wang, Z., Widgren, E. E., Sivashanmugam, P., O’Rand, M. G. and Richardson, R. T. (2005). Association of Eppin with Semenogelin on human spermatozoa. Biol. Reprod. 72(5), 1064–1070. doi:10.1095/biolreprod.104.036483

Young, S., Schiffer, C., Wagner, A., Patz, J., Potapenko, A., Herrmann, L., Nordhoff, V., Pock, T., Krallmann, C., Stallmeyer, B., Röpke, A., Kierzek, M., Biagioni, C., Wang, T., Haalck, L., Deuster, D., Hansen, J. N., Wachten, D., Risse, B., Behre, H. M., Schlatt, S., Kliesch, S., Tüttelmann, F., Brenker, C. and Strünker, T. (2024). Human fertilization in vivo and in vitro requires the CatSper channel to initiate sperm hyperactivation. J. Clin. Invest. 134(1). doi:10.1172/JCI173564

Zar, J. H. (2010). Biostatistical Analysis. New Jersey, USA: Pearson Prentice Hall.

Zeng, X.-H., Yang, C., Kim, S. T., Lingle, C. J. and Xia, X.-M. (2011). Deletion of the *Slo3* gene abolishes alkalization-activated K^+^ current in mouse spermatozoa. Proc. Natl. Acad. Sci. USA 108(14), 5879–5884. doi:10.1073/pnas.1100240108

Zeng, X.-H., Yang, C., Xia, X.-M., Liu, M. and Lingle, C. J. (2015). SLO3 auxiliary subunit LRRC52 controls gating of sperm KSPER currents and is critical for normal fertility. Proc. Natl. Acad. Sci. USA 112(8), 2599–2604. doi:10.1073/pnas.1423869112

Zhao, Y., Wang, H., Wiesehoefer, C., Shah, N. B., Reetz, E., Hwang, J. Y., Huang, X., Wang, T.-e., Lishko, P. V., Davies, K. M., Wennemuth, G., Nicastro, D. and Chung, J.-J. (2022). 3D structure and in situ arrangements of CatSper channel in the sperm flagellum. Nat. Commun. 13(1), 3439. doi:10.1038/s41467-022-31050-8

