## Supplementary Materials for "Semenogelin-1 Inhibition of Mouse Sperm Hyperactivation Reveals Two Functional Domains Modulating CatSper Channel"

### List of materials

#### Supplementary figures

**Figure S1.** SEMG1 immunolocalization in mouse epididymal spermatozoa.

**Figure S2.** Effects of mSEMG1 on mouse sperm motility.

**Figure S3.** Effects of mSEMG1 on sperm Em.

**Figure S4.** Effects of mSEMG1 and its truncated constructs on mouse sperm motility, $I_{CatSper}$, and viability.

**Figure S5.** Profile of the interaction between EPPIN and SEMG1.

**Figure S6.** Amido Black staining and Western blot analysis of recombinant mSEMG1 constructs.

**Figure S7**. Effect of dialyzed storage buffer on mouse sperm $I_{CatSper}$. .

#### Supplementary tables

**Table S1.** HS bath solution composition

**Table S2.** Pipette solution composition

**Table S3.** DVF bath solution composition

#### Supplementary videos

**Video S1.** Representative CASA videos displaying tracks of mouse spermatozoa incubated with mSEMG1.

**Video S2.** Representative CASA videos displaying tracks of mouse spermatozoa incubated with mSEMG1 truncated constructs.

### Supplementary figures

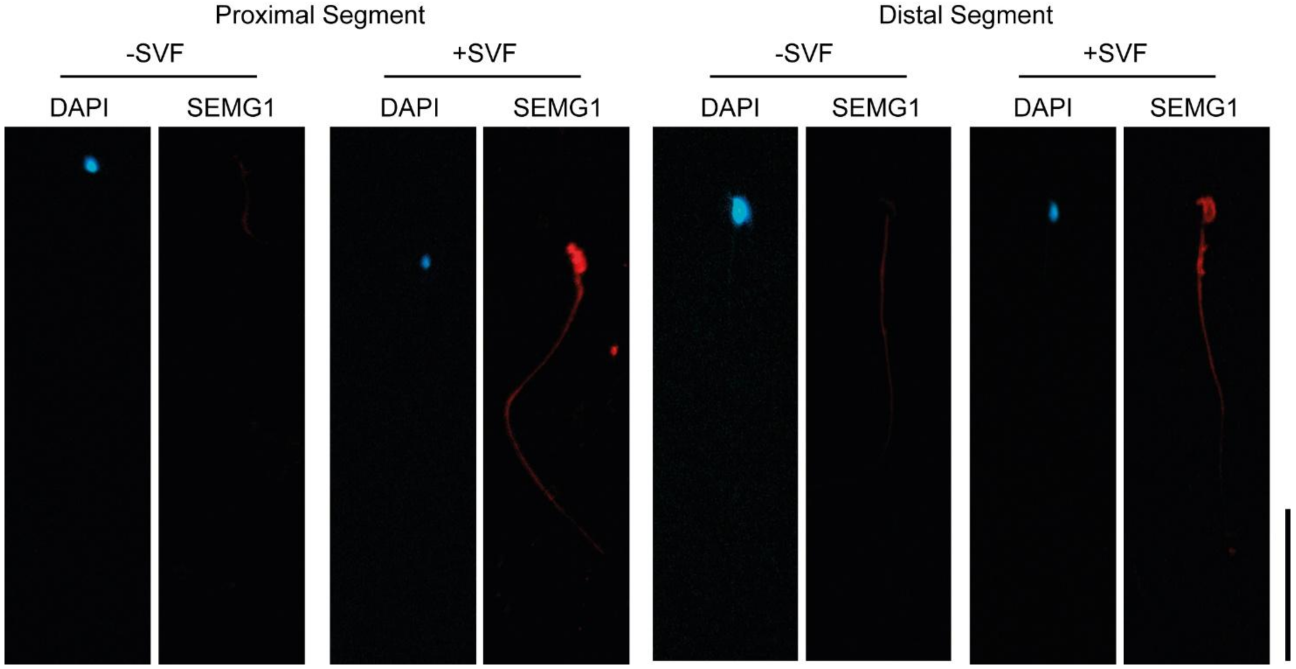


**Figure S1. SEMG1 immunolocalization in mouse epididymal spermatozoa.** Indirect immunolocalization of SEMG1 in spermatozoa collected from the proximal (initial segment and caput) and distal (corpus and cauda) regions of the mouse epididymis and incubated with (+SVF) or without (−SVF) seminal vesicle fluid. SEMG1 immunostaining was detected on the head and flagellum of mouse spermatozoa. Cell nuclei were counterstained with DAPI (blue). Scale bar: 50 µm. Results are representative of three independent experiments.

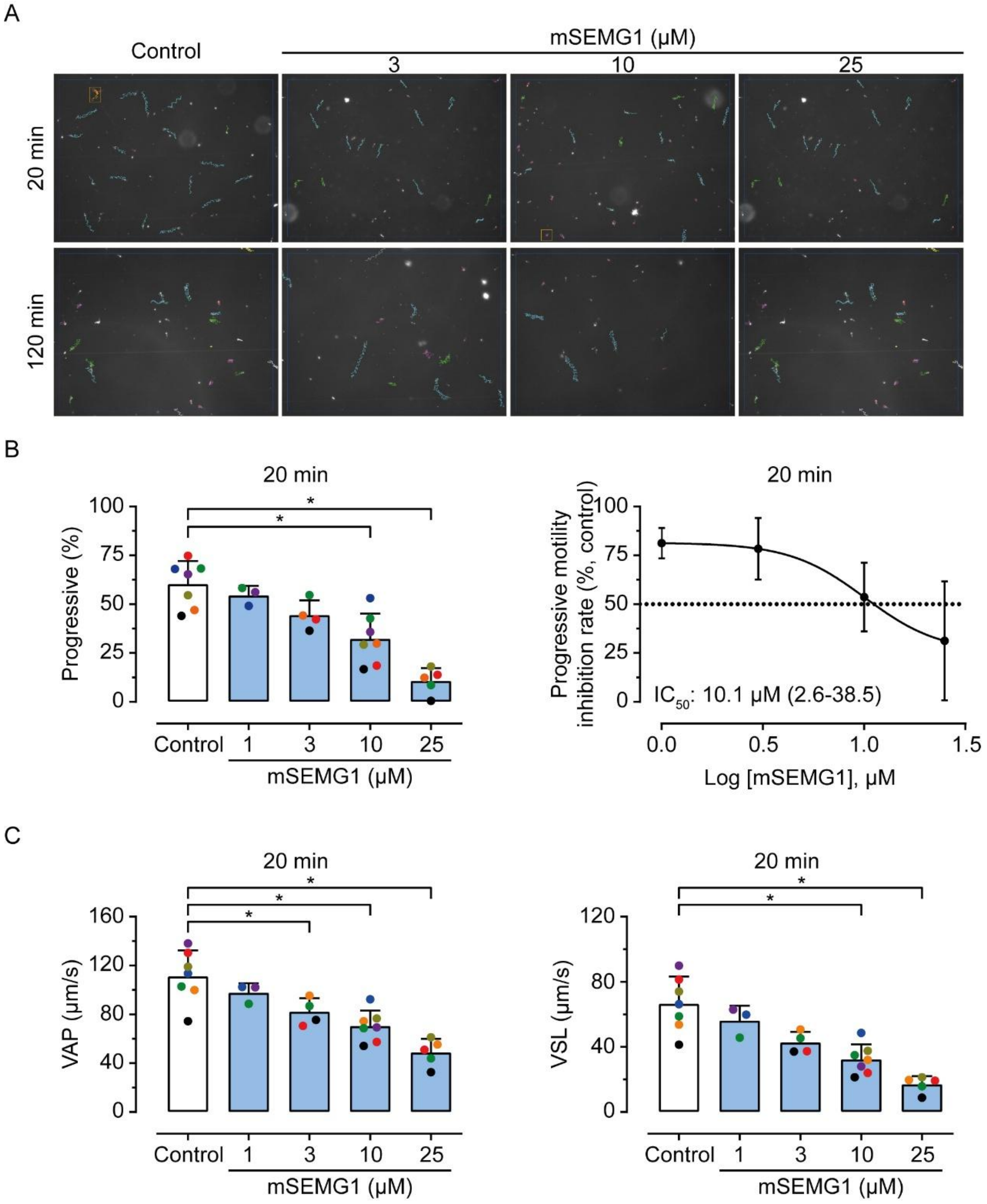


**Figure S2. Effects of mSEMG1 on mouse sperm motility.** (A) Representative CASA screenshots showing sperm tracks after incubation in capacitating HTF medium in the absence (control) or presence of increasing concentrations of mSEMG1 (1, 3, 10, and 25 µM) for 20 and 120 min. Green, motile cells; cyan, progressively motile cells; red, static cells. (B) Percentage of progressive motility (left) and concentration-response curve of progressive motility after 20 min of incubation (right). Concentration-response curve of progressive motility rate after incubation with mSEMG1. Data were expressed as mean percentage values relative to the control group, calculated as (% progressive spermatozoa in the treated sample divided by % progressive spermatozoa in the matched control) × 100. The half-maximal Inhibitory concentration ($IC_{50}$) and its 95% confidence interval) are presented. C) Average path velocity (VAP) and straight-line velocity (VSL) after 20 min of incubation. Dots of the same color represent values from the same independent experiment. Percentage data were transformed using the arcsine square-root transformation before statistical analysis. Asterisks indicate statistically significant differences from the control group ($p$ < 0.05, one-way ANOVA followed by Dunnett's multiple comparisons test). Results are presented as mean ± SD from independent experiments using sperm samples from 3–7 mice.

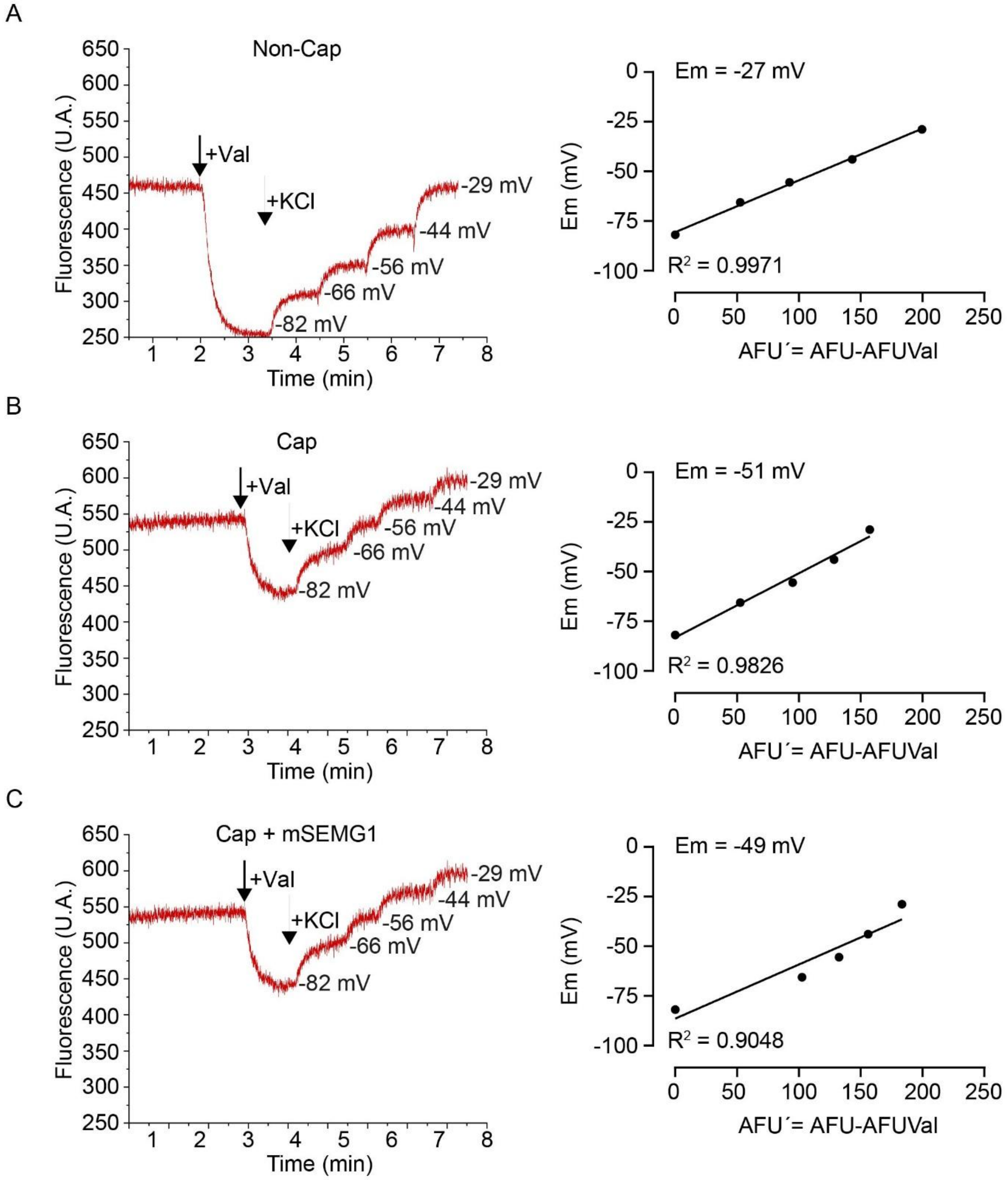


**Figure S3. Effects of mSEMG1 on sperm Em.** Representative Em recordings and corresponding calibration curves obtained under A) non-capacitating conditions (Non-Cap), B) capacitating conditions (Cap), and C) capacitating conditions in the presence of mSEMG1 (Cap + mSEMG1). Spermatozoa are loaded with DiSC3(5) (1µM) for ~3 min, transferred to a gently stirred quartz cuvette at 37°C, and the fluorescence is monitored with a spectrofluorometer at 620/670nm excitation/emission wavelength. Recordings began once a steady-state fluorescence signal was reached. Calibration was performed by adding valinomycin (1 µM, Val), followed by sequential additions of KCl. The Nernst equation was used to determine theoretical Em values corresponding to each arbitrary fluorescence unit (AFU) measured at KCl concentrations (points 1-4). To construct the calibration curve, the difference between each point and the fluorescence obtained after the addition of valinomycin (AFUVal) was calculated: AFU′(1′-4′). Em values were determined by linear interpolation of theoretical Em values plotted against initial AFU. Results represent mean ± SEM of specific signals from 3 experiments.

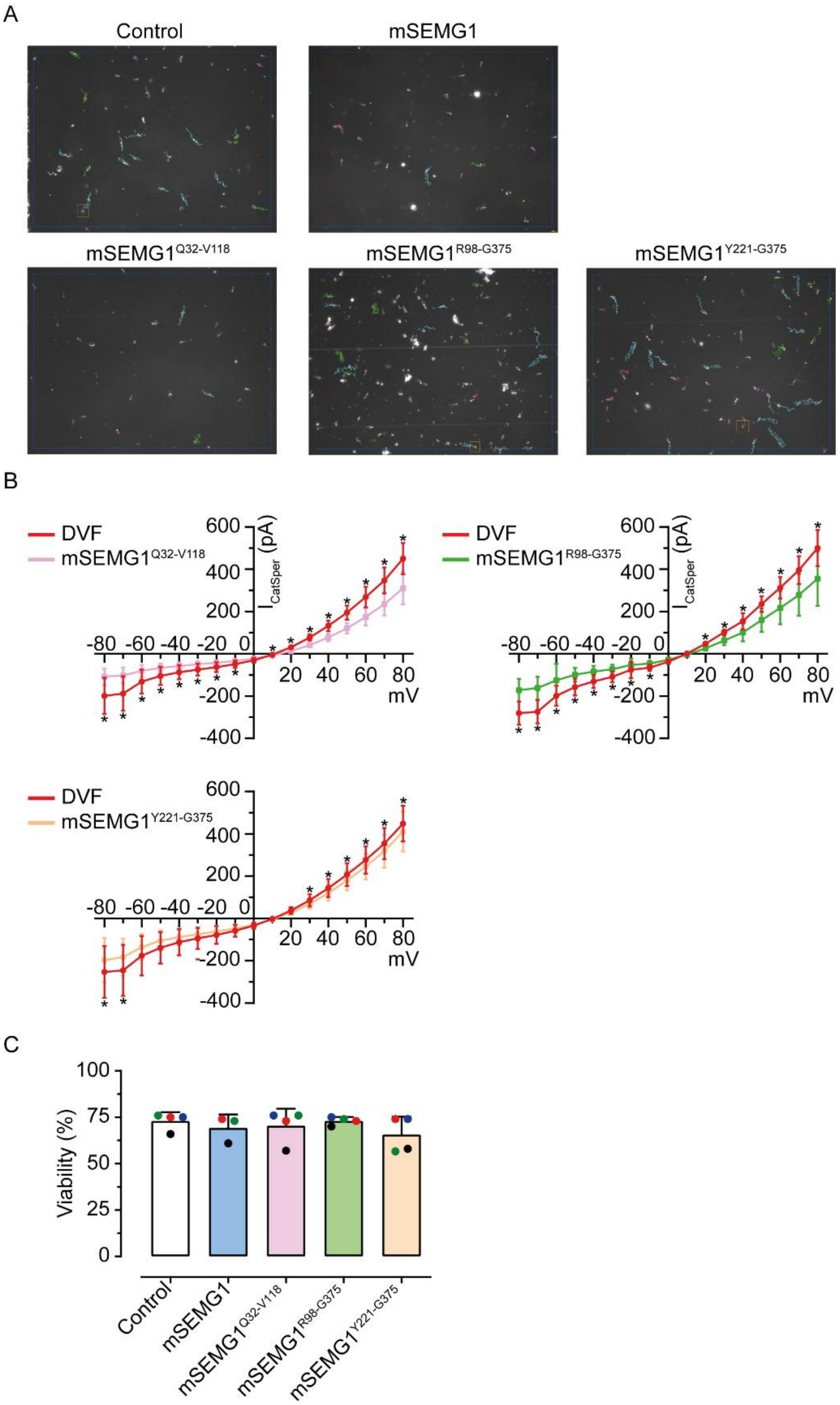


**Figure S4. Effects of mSEMG1 and its truncated constructs on mouse sperm motility, $I_{CatSper}$, and viability**. A) Representative CASA screenshots showing sperm tracks after incubation in capacitating HTF medium in the absence (control) or presence of mSEMG1 and its truncated constructs (25 μM) for 120 min. Green, motile cells; cyan, progressively motile cells; red, static cells. B) Current-voltage (I-V) relationships of CatSper were calculated from amplitudes shown in Figure 5 in the DVF solution before and after treatment with 5 μM mSEMG1 truncated constructs (mSEMG1$^{Q32–V118}$, mSEMG1$^{R98–G375}$, and mSEMG1$^{Y221–G375}$). Asterisks indicate statistically significant differences from the control group ($p < 0.05$, paired t-test). C) Percentage of viable spermatozoa after incubation with mSEMG1 truncated constructs (25 μM) for 120 min. Results are presented as mean ± SD from independent experiments using sperm samples from 3–7 mice.

A

```
1 mklsgfvsilvlfgllarvqgPSLADLLFPRRCPRFREECEHQERDLCTRDRDCPKKEKCCVFNCGKKCLNPQQ 74
                   qgPSLADLLFPRRCPRFREE (Q20E Ab)

75 DICSLPKDSGYCMAYFRRWWFNKENSTCQVFIYGGCQGNNNNFQSQSICQNACEKKSSLT 134
                            SMFVYGGCQGNNNNFQSKANC (S21C Ab)
```

B

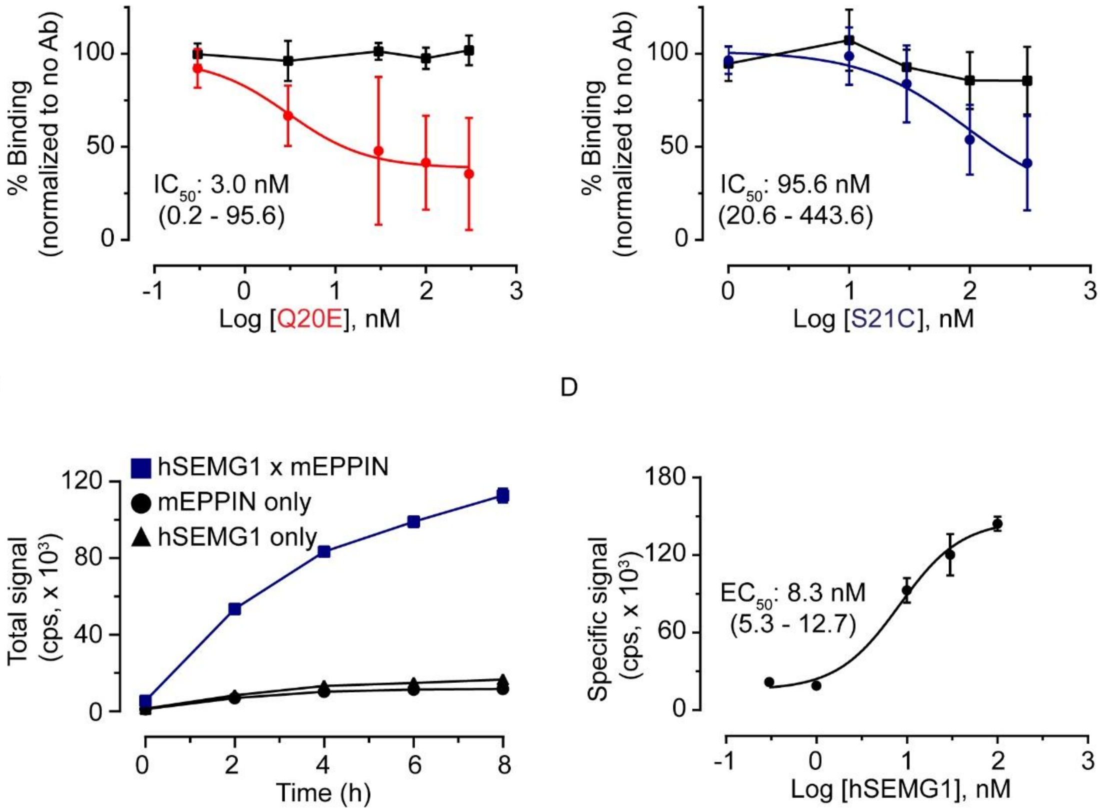


**Figure S5. Profile of the interaction between EPPIN and SEMG1.** (A) Primary sequence of mouse EPPIN indicating the epitopes recognized by anti-EPPIN antibodies Q20E and S21C. (B) Displacement of mSEMG1 from its binding sites on mEPPIN by Q20E or S21C antibodies. mEPPIN was incubated with mSEMG1 in the presence of increasing concentrations of each antibody. Specific signal at each data point was determined by subtracting the background from the total signal and normalized to the specific signal in the absence of antibody. Control experiments using AlphaScreen TruHits (black) were performed in the presence of Q20E or S21C under similar conditions. Results are expressed as mean ± SD of specific signals from 5-7 experiments. (C) Interaction of mEPPIN with human recombinant SEMG1 (hSEMG1). mEPPIN was incubated with hSEMG1 in a time-course experiment. Background signal was detected when beads were incubated in the absence of mEPPIN or mSEMG1. The graph shows a representative experiment from two independent experiments; each performed in triplicate. (D) Concentration-response curve for hSEMG1 with a constant concentration of mEPPIN (10 nM). Right: Concentration-response curve for mEPPIN with a constant concentration of mSEMG1 (30 nM). A specific signal for each data point was determined by subtracting the background signal from the total signal. Results represent the mean ± SD of specific signals from 2 experiments. Cps = counts per second.

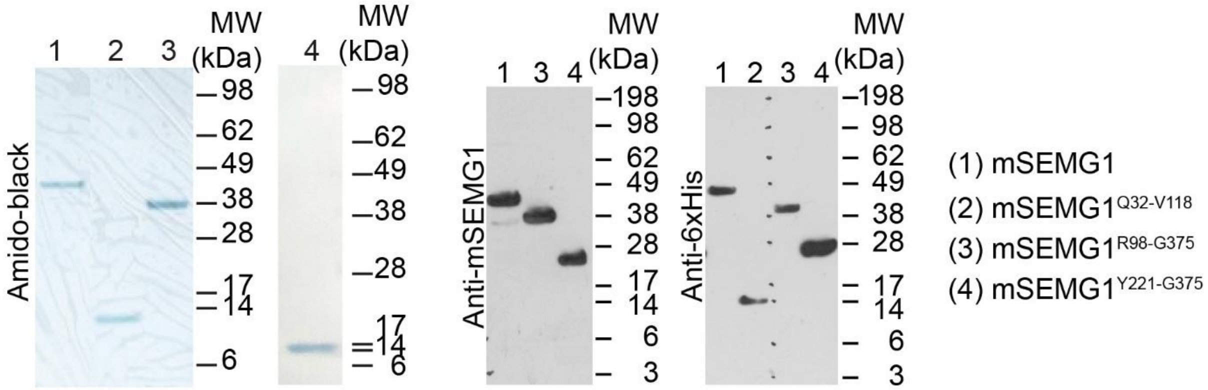


**Figure S6. Amido Black staining and Western blot analysis of recombinant mSEMG1 constructs.** (Left) Representative Amido Black-stained membrane showing the purified recombinant mSEMG1 constructs (0.5–1 μg) used in this study. A single predominant band was observed for each purified protein, supporting the purity of the protein preparations used in the functional assays. Lane 1: 6×His-mSEMG1 (Q32–G375), ~40 kDa; Lane 2: 6×His-mSEMG1$^{Q32–V118}$ (Q32–V118), ~11 kDa; Lane 3: 6×His-mSEMG1$^{R98–G375}$ (R98–G375), ~31 kDa; Lane 4: 6×His-mSEMG1$^{Y221–G375}$ (Y221–G375), ~19 kDa. Molecular weight markers (kDa) are shown on the right. (Right) The identity and purity of all recombinant proteins was independently confirmed by Western blot using anti-SEMG1 and anti-6×His antibodies. The Western blot was performed using a separate gel from that shown for the Amido Black staining.

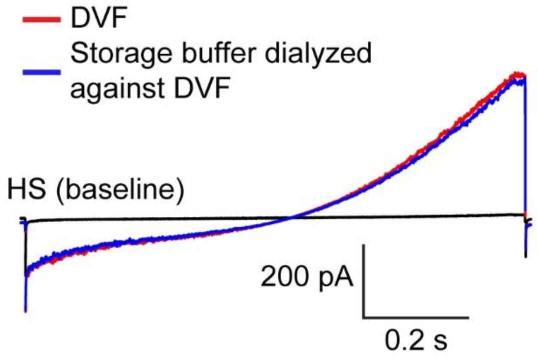


**Figure S7**. **Effect of dialyzed storage buffer on mouse sperm $I_{CatSper}$.** Mouse sperm $I_{CatSper}$ recording in response to a voltage ramp in DVF before (red) and after treatment with protein storage buffer dialyzed against DVF (blue). Baseline currents are recorded in HS (black). All traces were obtained from the same spermatozoon. The result represents an experiment conducted with one spermatozoon.

### Supplementary tables

**Table S1.** HS bath solution composition

| Chemicals | mM |
|---|---|
| **NaCl** | 135 |
| **KCl** | 5 |
| **CaCl2 x 2 H2O** | 2 |
| **MgSO4 x 7 H2O** | 1 |
| **HEPES** | 20 |
| **Glucose** | 5 |
| **Sodium lactate (60% w/w)** | 10 |
| **Sodium pyruvate** | 1 |

**Table S2.** Pipette solution composition

| Chemicals | mM |
|---|---|
| **CsMeSO3** | 130 |
| **HEPES** | 70 |
| **EDTA** | 2 |
| **EGTA** | 3 |
| **CsCl** | 1 |

**Table S3.** DVF bath solution composition

| Chemicals | mM |
|---|---|
| **CsMeSO3** | 140 |
| **HEPES** | 40 |
| **EDTA** | 1 |
